# Chemical signatures of human odour generate a unique neural code in the brain of *Aedes aegypti* mosquitoes

**DOI:** 10.1101/2020.11.01.363861

**Authors:** Zhilei Zhao, Jessica L. Zung, Alexis L. Kriete, Azwad Iqbal, Meg A. Younger, Benjamin J. Matthews, Dorit Merhof, Stephan Thiberge, Martin Strauch, Carolyn S. McBride

## Abstract

A globally invasive form of the mosquito *Aedes aegypti* specializes in biting humans, making it an efficient vector of dengue, yellow fever, Zika, and chikungunya viruses. Host-seeking females strongly prefer human odour over the odour of non-human animals, but exactly how they distinguish the two is not known. Vertebrate odours are complex blends of volatile chemicals with many shared components, making discrimination an interesting sensory coding challenge. Here we show that human and animal odour blends evoke activity in unique combinations of olfactory glomeruli within the *Aedes aegypti* antennal lobe. Human blends consistently activate a ‘universal’ glomerulus, which is equally responsive to diverse animal and nectar-related blends, and a more selective ‘human-sensitive’ glomerulus. This dual signal robustly distinguishes humans from animals across concentrations, individual humans, and diverse animal species. Remarkably, the human-sensitive glomerulus is narrowly tuned to the long-chain aldehydes decanal and undecanal, which we show are consistently enriched in (though not specific to) human odour and which likely originate from unique human skin lipids. We propose a model of host-odour coding wherein normalization of activity in the human-sensitive glomerulus by that in the broadly-tuned universal glomerulus generates a robust discriminatory signal of the relative concentration of long-chain aldehydes in a host odour blend. Our work demonstrates how animal brains may distil complex odour stimuli of innate biological relevance into simple neural codes and reveals novel targets for the design of next-generation mosquito-control strategies.

---

The discrimination of odour cues is a challenging problem faced by animals in nature. Decades of olfactory research has revealed the principles by which animals may identify individual compounds or simple mixtures – using combinatorial codes for flexible, learned behaviours^1–6^ or labelled lines for hard-wired, innate responses^7–12^. However, most natural odours are blends of tens to hundreds of compounds^13–15^. How animals evolve to efficiently recognize these more complex stimuli, especially those with important innate meaning, is poorly understood^16–19^.

This problem is particularly relevant for *Aedes aegypti* mosquitoes. A globally invasive subspecies of *Ae. aegypti* has recently evolved to specialize in biting humans and thus become the primary worldwide vector of human arboviral disease^20,21^. Females rely heavily on their sense of smell to choose among potential hosts^22^ and show a robust preference for human odour over the odour of non-human animals^21,23,24^ (hereafter ‘animals’) (Fig. 1a–d). The apparent ease with which they distinguish these stimuli is remarkable since vertebrate odours are complex blends of relatively common compounds that are frequently shared across species^13,25–27^. Females require a multi-component blend for strong attraction^28–30^ and may discriminate based on the ratios in which different components are mixed^31,32^. Understanding exactly which features of human odour are used for discrimination and how these features are represented at the neural level would provide basic insight into olfactory coding and potential targets for use in vector control.

**Fig. 1.**
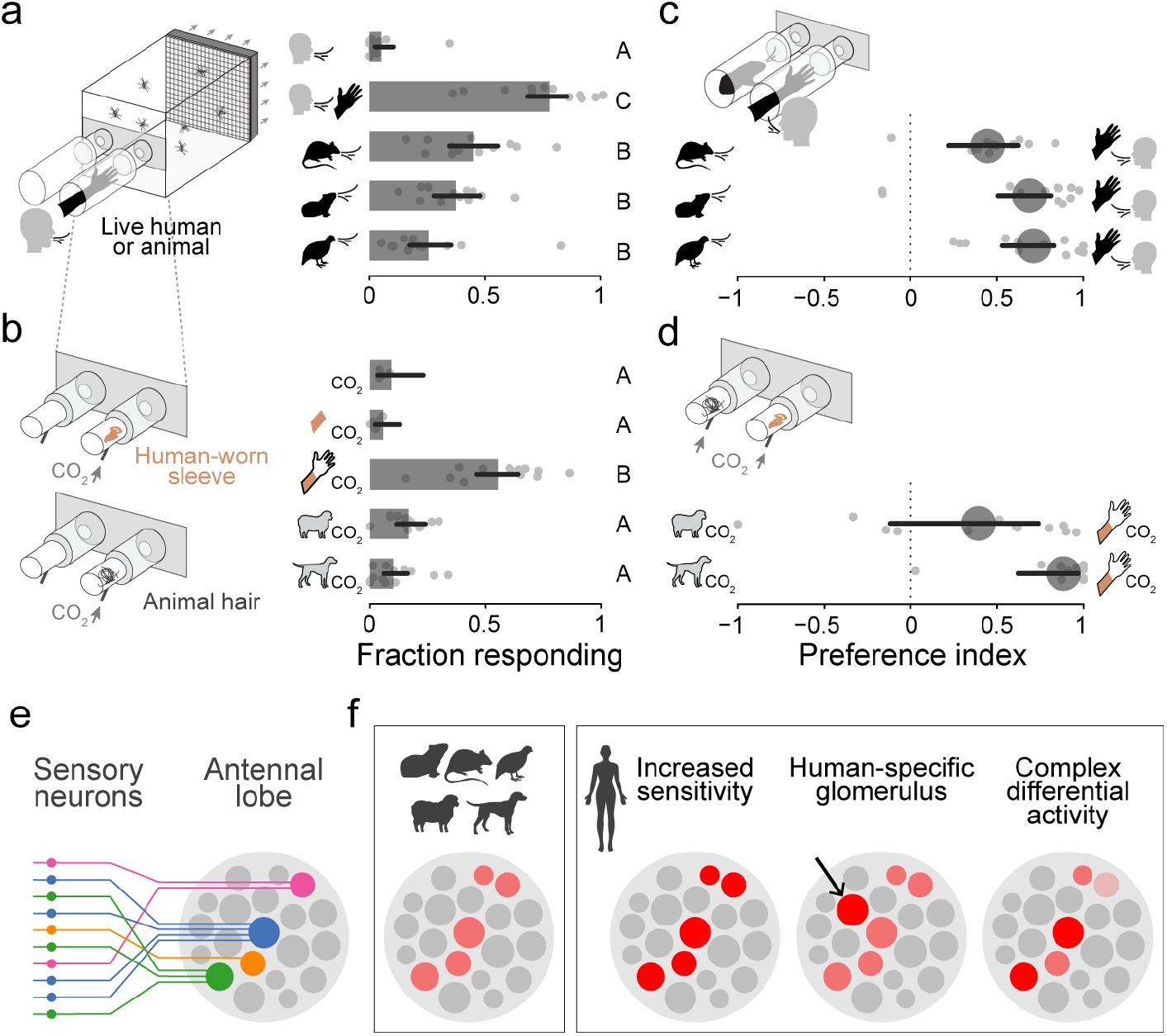
Preference of *Aedes aegypti* mosquitoes for human odour and possible coding mechanisms. **a–d**, Response of female *Ae. aegypti* mosquitoes to human and animal odours in no-choice (**a,b**) and choice (**c,d**) olfactometer trials. Bars (or circles) and lines represent means and 95% confidence intervals from beta-binomial mixed models (n=12 trials/comparison evenly spread across 6 humans, 2 rats, 2 guinea pigs, 1 quail, wool from 1 sheep, and hair from 4 dogs). Response to exhaled human breath (**a**, top), synthetic CO_2_ (**b**, top), or unworn control sleeves (**b**, second from top) was minimal in the absence of human or animal odour. **e**, All olfactory sensory neurons that express the same receptor complex (same colour) send axons to a single glomerulus in the antennal lobe. **f**, Schematics show different ways in which the neural activity evoked by human and animal odours in the antennal lobe may differ, allowing mosquitoes to discriminate. Shades of red indicate different levels of neural activity.

Mosquitoes detect volatile chemical cues using thousands of olfactory sensory neurons scattered across the antennae and maxillary palps^33^. Carbon dioxide is detected by a single class of neurons that express a gustatory receptor complex^34–37^, while acids and amines are detected by multiple neural classes that express receptors in the ionotropic receptor (IR) family^38–41^. These odorants and neurons are critical for baseline host attraction in *Ae. aegypti*^28,36,42–44^. However, for fine-grained discrimination among host species, females rely heavily on a more diverse set of ligands recognized by neurons that express receptors in the odorant receptor (OR) family^24,45^. Females carrying mutations in the conserved OR co-receptor *orco* are attracted to hosts, but fail to discriminate strongly between humans and animals^45^. In the brain, all neurons that express the same ligand-specific receptor complex send axons to a single olfactory glomerulus within the antennal lobe^46^ (Fig. 1e), making it an ideal location to decipher the coding of complex host odour blends across sensory neuron classes^3,18^. Here we set out to determine how representations of human and animal odours differ at this critical junction (Fig. 1f) and what chemical features underlie this difference.

## Novel reagents and methods for olfactory imaging in mosquitoes

We first developed tools to visualize odour-evoked responses in olfactory sensory neurons where they enter antennal lobe glomeruli. Focusing on the subset of neurons in the OR pathway, we used CRISPR/Cas9 to generate knock-in mosquitoes that express the calcium indicator GCaMP6f under the endogenous control of the *orco* locus^47–50^ (Fig. 2a). Transgenic individuals showed GCaMP6f expression in sensory neurons on the adult antenna and maxillary palp that project to approximately 34 of 60 glomeruli in the dorso-medial antennal lobe (Fig. 2b, Extended Data Fig. 1). We also observed GCaMP6f in sensory neurons that project to the subesophageal zone (SEZ) from the labellum^51^ and, most likely, the legs^52^ (Extended Data Fig. 1j–l). Together with a two-photon microscope custom-designed for fast, volumetric imaging (Fig. 2c) and a novel analytical pipeline (Fig. 2d, Extended Data Fig. 2), the new strain allowed us to capture odour-evoked responses in all Orco+ glomeruli at ~4 Hz.

**Fig. 2.**
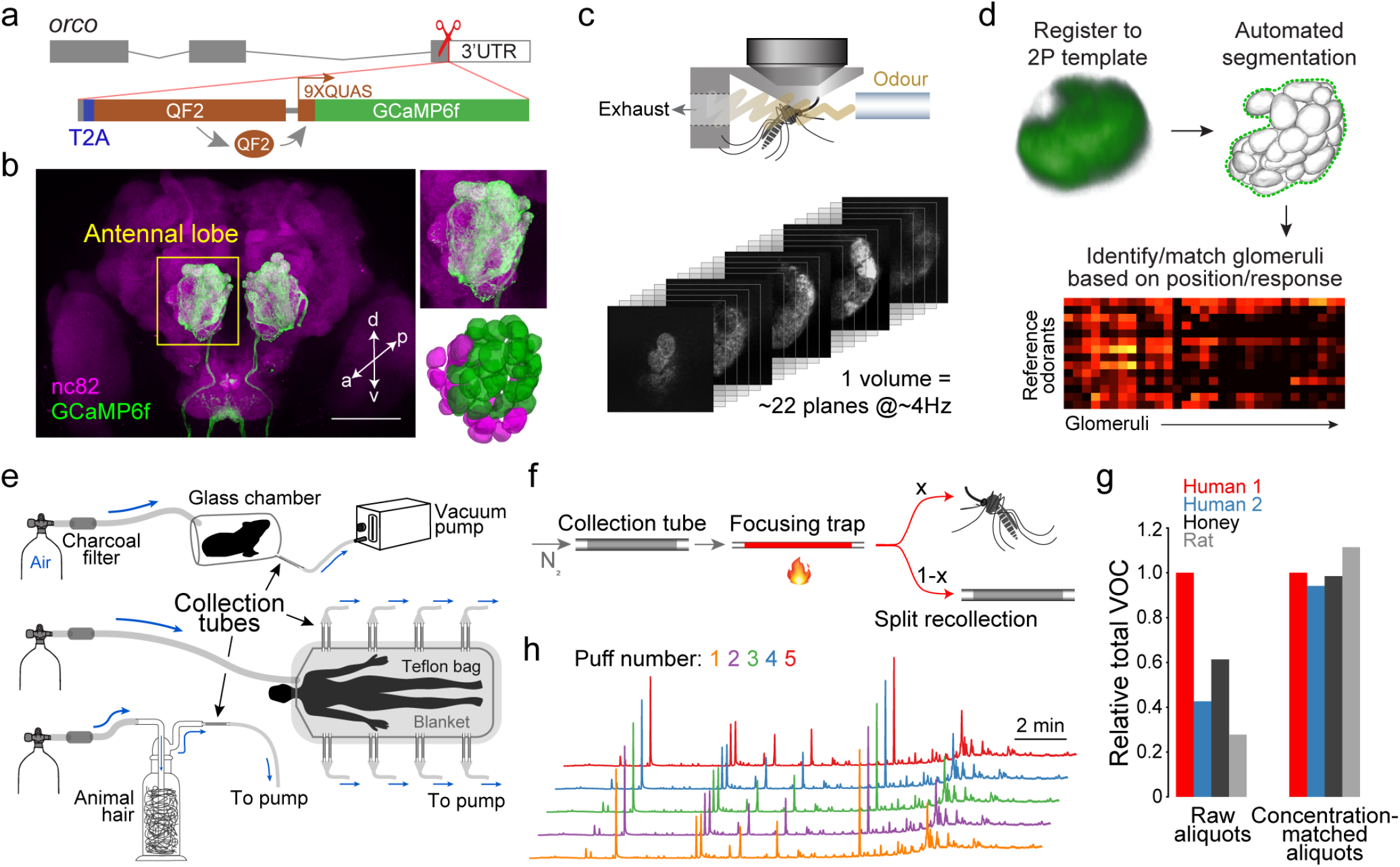
Novel reagents and methods for imaging *Ae. aegypti* olfactory circuits. **a,** Gene-targeting strategy used to drive GCaMP6f expression in Orco+ sensory neurons while preserving native *orco* function. **b**, Antibody staining of *orco-T2A-QF2-QUAS-GCaMP6f* adult female brain, with close-up of antennal lobe (upper right inset). Lower right inset shows 3D reconstruction with approximately 34 Orco+ (green) and 20 Orco- (magenta) glomeruli. Scale bar, 100μm. **c**, Schematic of mosquito prep and stack of movies from fast volumetric imaging. **d**, Novel analysis pipeline. The final glomerulus-matching step can be completed manually or using an automated algorithm (see Methods, Extended Data Fig. 2). **e**, Odour sampling set-ups for live animals/milkweed (top), humans (middle), and animal hair/honey (bottom). **f**, Schematic of two-stage thermal desorption for delivery of complex odour samples. Extracts are transferred from the collection tube to a sorbent-filled focusing trap via slow heating and nitrogen flow. The focusing trap is then heated ballistically (to 220° C in ~3 sec) to release all odorants, which cool to room temperature before reaching the mosquito in the form of a 3–4 sec puff (see Extended Data Fig. 3). The final odour stream is split such that an adjustable percentage flows to the mosquito, while the remainder can be recollected. **g**, Verification of the concentration-matching procedure for four representative odour samples (see Methods, Extended Data Fig. 3). Volatile organic content (VOC) was measured via GC-MS before (left) and after (right) matching. **h**, GC-MS chromatograms of 5 consecutive puffs of the same human sample demonstrating consistency of blend ratios and absolute abundance. Arbitrary y-axis units not shown.

We next collected natural odours and developed methods to faithfully deliver these stimuli to mosquitoes during imaging. We sampled odour from humans (n=8), rats (n=2), guinea pigs (n=2), quail (n=2), sheep wool (n=1), dog hair (n=4), and two nectar-related stimuli that mosquitoes find attractive – milkweed flowers^53^ and honey^45^ (Fig. 2e). Individual human samples were kept separate, while those from animals were pooled by species. For delivery, stored odour extracts are usually eluted from sorbent collection tubes with a solvent and then allowed to evaporate from a vial, septum, or filter paper^17,54,55^. However, the diverse odorants in a blend often require different solvents and will evaporate from solution at different rates based on volatility^56,57^, changing the character of a blend over time. We therefore developed a novel odour-delivery system involving thermal desorption^58^ that allowed us to deliver natural odour extracts directly from sorbent tubes to mosquitoes with precise quantitative control (Fig. 2f, Extended Data Fig. 3). Importantly, we were able to match the total odour concentration of diverse samples delivered to the same mosquito (Fig. 2g) and to deliver replicate puffs of the same sample to different mosquitoes, all while maintaining the original blend ratios (Fig. 2h).

## Human and animal odours activate unique combinations of antennal lobe glomeruli

With these new tools and odour samples in hand, we set out to characterize the response of Orco+ glomeruli to host odours. There are several ways in which the activity of these glomeruli might allow female mosquitoes to discriminate human and animal blends (Fig. 1f). A single set of host-responsive glomeruli may simply be more sensitive to human blends than to animal blends. Alternatively, one or more glomeruli may be exclusively activated by either human or animal blends. Third, a more complex pattern of differential activity in common glomeruli may be required for discrimination.

To explore these possibilities, we characterized antennal lobe responses to the odour of a single human at concentrations ranging from 1/25X to 5X, where 1X matches the concentration exiting the collection bag during a representative human odour extraction (Fig. 2e, see Methods). The number of responding glomeruli increased with dose, with only two glomeruli responding consistently at the middle doses (cyan and green arrows) and two to five more joining the ensemble at the highest dose (Fig. 3a,b). Although there may be additional responses below the sensitivity threshold of our preparation, we were struck by the sparseness of this activity.

**Fig. 3.**
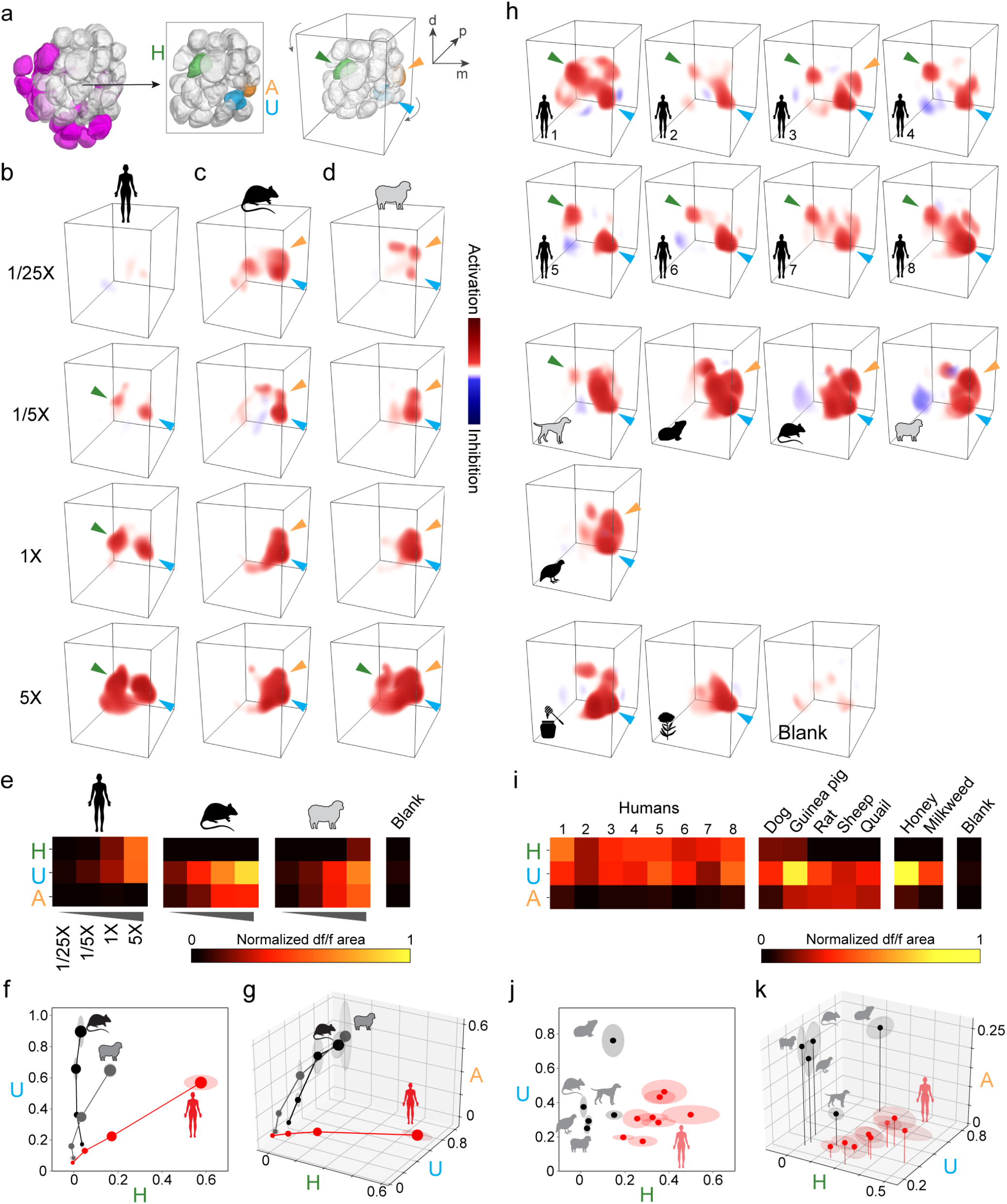
Human and animal odours activate unique combinations of antennal lobe glomeruli. **a,** Antennal lobe reconstructions highlighting Orco+ glomeruli imaged here (left, grey), three glomeruli that dominate host odour representations (middle, with a few anterior glomeruli removed to reveal the posterior U and A glomeruli), and the angle from which they are viewed in 3D renderings (right). U, universal; H, human-sensitive; A, animal-sensitive. **b–d**, 3D renderings of the response of a single representative female mosquito to human, rat, and sheep odour. Arrowheads indicate focal glomeruli from (**a**). **e–g**, Mean activity evoked by stimuli in (**b–d**) across n=4 mosquitoes, visualized as a heat map (**e**) or the relative activation of individual glomeruli (**f,g**; dot size reflects dose, shading around dots indicates SEM). **h–k**, Same as (**b–g**) but showing response to the odour of 8 individual humans, 5 animal species, and 2 nectar stimuli all at 1X total odour concentration. n=5 mosquitoes for (**i-k**). Human subject numbers in (**h,i**) correspond to those in Fig. 4a,b.

We next examined the pattern of activity evoked by two animal odours at the same four concentrations. We chose rat and sheep because they are common in human environments and provided ample odour in our extractions. One of the two glomeruli most sensitive to human odour was also highly sensitive to the animal odours (Fig. 3c,d, cyan arrow), while the second was insensitive to rat and responded to only the highest dose of sheep (Fig. 3c,d, green arrow). Since we cannot reliably map these glomeruli to existing antennal lobe atlases^59,60^ without molecular markers, we call these the ‘universal’ (U) and ‘human-sensitive’ (H) glomeruli, respectively (Fig. 3a). We call a third glomerulus the ‘animal-sensitive’ (A) glomerulus, because it was strongly activated by both rat and sheep but not human (Fig. 3a–d, orange arrow). Remarkably, the relative activity of just these two or three glomeruli cleanly separated human and animal odours across the concentration gradient (Fig. 3e–g).

The behavioural preference of *Ae. aegypti* for human odour is robust to variation among individual humans and across animal species (Fig. 1a–d). We therefore asked whether the patterns of glomerular activity described above were similarly robust by imaging responses to odour from 7 additional humans (8 total), 3 more animal species (5 total), and the two nectar-related stimuli at a single concentration (1X). The U glomerulus was again strongly activated by all odour extracts, including the two nectar odours, while H and A glomeruli were most strongly activated by human and animal odours, respectively (Fig. 3h,i, Extended Data Fig. 4). Taken together, we again saw clean separation of humans and animals in the neural space of just these two or three glomeruli (Fig. 3j,k).

While U, H, and A dominated host-odour responses in our experiments, we wanted to make sure we had not missed additional discriminatory signals. We therefore used an automated pipeline to match as many glomeruli as possible across mosquito brains (Extended Data Fig. 2c). In a principal components analysis, U, H, and A again explained most of the observed variation, with H and A loaded most strongly on the principal component separating humans from animals (Extended Data Fig. 5). A fourth glomerulus just posterior to U also responded to vertebrate odours and may be the target of well-known, 1-octen-3-ol-sensing neurons that project to this region from the palp^59,61^. In summary, our results indicate that human and animal odours activate unique combinations of glomeruli in the antennal lobe of *Ae. aegypti*, generating a discriminatory neural signal that is robust to both concentration and individual/species variation.

## Human odour is enriched for select ketones and long-chain aldehydes

The neural response to human odour must be traceable to chemical features of human odour blends. Human blends contain an array of common volatile compounds that originate from skin secretions, the skin microbiome, or their interaction^13^. They differ consistently from animal blends in the relative abundance of at least two or three components, but quantitative, cross-species comparisons are rare and almost always focus on a single compound^24,26,27,62,63^. We therefore lack a clear, comprehensive picture of the relative ratios and other chemical features mosquitoes may use to discriminate.

To fill this gap, we analysed the composition of the human, animal, and nectar-related odour samples used for imaging, plus 8 new human samples (Fig. 4a, Extended Data Fig. 6a–d). Importantly, we quantified the abundance of all compounds that made up at least 2% of any blend (by GC-MS peak area), excluding acids (sensed primarily by the IR pathway) and other highly polar or volatile compounds that cannot be quantified reliably within the same framework (see Methods). Consistent with previous work, the vertebrate odours were dominated by aliphatic aldehydes^13,25,26^, whereas nectar odours were enriched for terpenes^14^ (Fig. 4a). Also as expected, human and animal odours shared almost all of the same components (Extended Data Fig. 6c), highlighting the challenge mosquitoes face: the presence of specific compounds is by itself insufficient for discrimination.

**Fig. 4.**
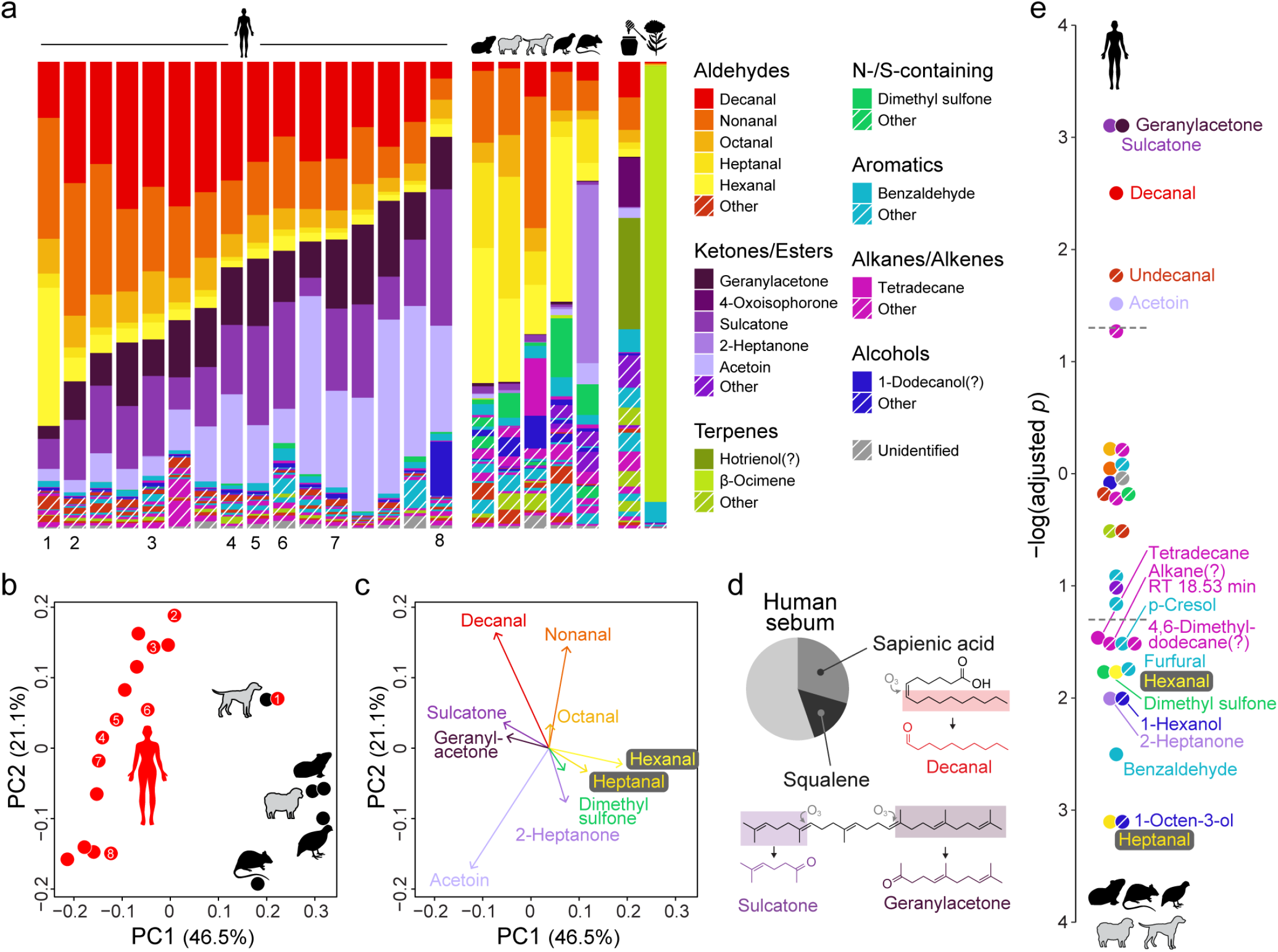
Human and animal odour blends differ in the relative concentration of key compounds. **a**, Odour profiles showing relative abundance of compounds that made up >2% of at least one sample. Named compounds made up >10% of at least one sample or an average of >1% across samples. Question marks indicate tentative identifications (see Methods). Animal samples pooled by species prior to analysis (n=4 dogs, 2 guinea pigs, 1 sheep, 2 rats, 2 quail). Human samples used for imaging (Fig. 3h,i) are numbered. **b**, Unscaled principal components analysis on human and animal data from (**a**). **c**, Top ten loadings on first two principal components from (**b**). **d**, Proportion of human sebum made up of sapienic acid and squalene^81^. Oxidation of the two lipids produces volatile compounds enriched in human odour^64^. **e**, *P*-values from Kolmogorov-Smirnov tests for a difference in the relative abundance of each odorant between humans and animals (corrected for multiple testing using the Benjamini-Hochberg procedure). Values extend up or down from zero for human- or animal-biased odorants, respectively. Dashed lines mark *P*=0.05.

Despite the overlap in blend components, human and animal samples differed consistently in blend ratios, leading to clear separation in a principal components analysis (PCA) (Fig. 4b, Extended Data Fig. 6e). Loadings on the human–animal axis of the PCA showed that human odour was enriched for the ketones sulcatone and geranylacetone (Fig. 4a,c). Another striking feature of human odour was the high relative abundance of long-chain aldehydes and low relative abundance of short-chain aldehydes: human odour had more decanal (10 carbons) and less hexanal and heptanal (6 and 7 carbons) than the animal odours (Fig. 4a,c). While the two ketones and decanal are widely recognized as abundant in human odour^13^, consistent enrichment compared to animal odours has only been previously documented for sulcatone^24^. Interestingly, these three compounds are oxidation products of squalene and sapienic acid^64^, unique components of human sebum that may play a role in skin protection^65–67^ (Fig. 4d).

The unscaled PCA gives the most weight to abundant compounds, but mosquitoes may also be sensitive to minor components^68,69^. We therefore carried out additional, compound-specific comparisons of relative abundance, revealing several more differences (Fig. 4e, Extended Data Fig. 6f). Most notably, human odour was significantly enriched not only for the two ketones and decanal, but also for a second long-chain aldehyde (undecanal, 11 carbons) and acetoin. Human odour can thus be distinguished from animal odours by the relative abundance of a diverse set of compounds, none of which signify ‘human’ in isolation but which come together in characteristic ratios to produce a uniquely human bouquet.

## The human-sensitive glomerulus is narrowly tuned to long-chain aldehydes

To connect the unique pattern of neural activity evoked by human odour (Fig. 3) to its chemical composition (Fig. 4), we conducted additional imaging with synthetic odorants and blends delivered using standard approaches (Fig. 5a, Extended Data Fig. 7). We first asked if the neural response to a representative human sample could be explained by the response to its major components delivered either individually or in a ‘combo’ blend. We considered each of the 11 most abundant compounds in the human sample with two exceptions: geranylacetone was excluded because it is unstable under lab conditions, and acetoin was delivered singly but absent from the combo since it requires a different solvent. We directly measured delivered stimuli using GC-MS and individually calibrated the liquid dilution ratios to replicate vapour-phase concentrations in the human sample at 1X (Fig. 5a,b)^70^.

**Fig. 5.**
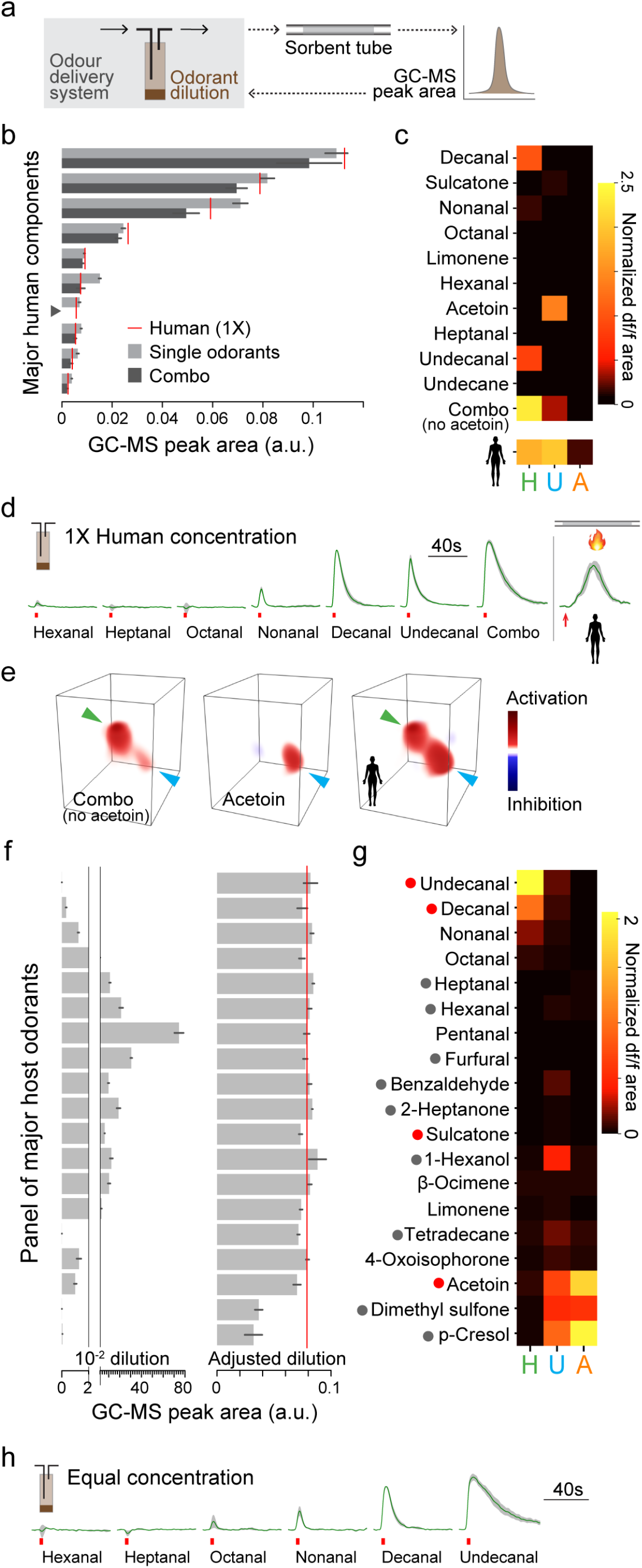
Tuning of focal glomeruli to major host odorants can explain response to blends. **a,** Single-odorant delivery system and procedure used to estimate and adjust vapour-phase stimulus concentrations. **b**, GC-MS peak area of 3-sec puffs of major human odorants and their mixture (combo) when diluted to generate the same vapour-phase concentrations found in 1X human odour (red lines). Odorant names same as in (**c**). Acetoin was excluded from the mixture (grey arrowhead) because it is not soluble in paraffin oil. n=4–5 puffs. **c**, Mean normalized response of 3 focal glomeruli to stimuli from (**b**). n=4 mosquitoes. **d**, Time traces for response of the H glomerulus to straight-chain saturated aldehydes, the combo from (**b,c**), and 1X human odour delivered by thermal desorption. **e**, 3D rendering of the antennal lobe response to the combo, acetoin, and 1X human odour in one representative mosquito. Coloured arrows point to focal glomeruli (green: H; cyan: U). **f**, GC-MS peak area of 3-sec puffs of major host odorants at a standard 10^-2^ v/v liquid dilution (left) and a second dilution individually adjusted to equalize vapour-phase concentration (right). Red line shows target concentration (set equal to sulcatone in 1X human odour). n=3–9 puffs. Odorant names same as in (**g**). p-Cresol and dimethyl sulfone were too insoluble/nonvolatile to achieve the desired concentration. **g**, Mean normalized response of 3 focal glomeruli to adjusted, equal vapour-phase stimuli from (**f**). Dots before names indicate human-biased (red) and animal-biased (grey) compounds. n=4–5 mosquitoes. **h**, Time traces for response of the H glomerulus to straight-chain, saturated aldehydes from (**f,g**). Bars/black lines in (**b,f**) and green lines/grey shading in (**d,h**) indicate mean ± SEM.

Decanal, undecanal, and the combo stimulus that contained them all evoked strong and prolonged activity in H (Fig. 5c–e, Extended Data Fig. 8a). The U glomerulus was strongly activated by acetoin and weakly activated by the combo of non-acetoin compounds (Fig. 5c,e), possibly the sum of a number of tiny individual responses (Extended Data Fig. 8a). No human odour components evoked activity in the A glomerulus. Genetic mapping previously implicated a sulcatone receptor in preference evolution in *Ae. aegypti*^24^. While we did not see consistent activity in response to this compound at its concentration in 1X human odour (Fig. 5c), several glomeruli responded at higher doses (data not shown), suggesting it may be important at close range. Taken together, the response to 1X human odour is largely explained by individual responses to a subset of perceptually dominant components, including long-chain aldehydes and acetoin (Fig. 5e).

The strong response of the H glomerulus to physiological concentrations of decanal and undecanal in human odour, but not the shorter-chain compounds (Fig. 5d), suggests it may be selectively tuned to long-chain aldehydes. To rigorously test this hypothesis and more broadly explore the tuning of all three focal glomeruli, we next imaged the response of H, U, and A to a panel of 19 compounds all delivered at the same concentration (Fig. 5f,g). The panel included seven straight-chain, saturated aldehydes and most other odorants that made up >3% of any vertebrate or nectar-related blend in our collection (see Methods). Since compound-specific volatilities varied across two to three orders of magnitude, we again directly measured delivered stimuli and individually calibrated v/v dilution ratios (Fig. 5a) to equalize vapour-phase concentrations (Fig. 5f, target concentration set to that of sulcatone in 1X human odour).

As hypothesized, the H glomerulus responded selectively to long-chain aldehydes (Fig. 5g,h). Both response amplitude and duration increased with aldehyde chain length, from the 6-carbon hexanal that evoked no response to the 11-carbon undecanal that evoked strong activity lasting 40+ seconds beyond the 3-second puff (Fig. 5h). This narrow tuning explains why H was most strongly activated by human blends, which are enriched for long-chain aldehydes. The U glomerulus, in contrast, showed surprisingly broad tuning. It responded to 10 of 19 compounds in the panel, including human-biased, animal-biased, and unbiased odorants (Fig. 5g, Extended Data Fig. 8b), consistent with its universal response to complex blends. The A glomerulus was strongly activated by two animal-biased compounds previously shown to be enriched in vertebrate faeces and urine (dimethyl sulfone^71^ and p-cresol^15^) as well as the higher-than-human dose of acetoin used in this experiment (~10 times that in 1X human odour) (Fig. 5g, Extended Data Fig. 8b). Since the smaller animal species in our sample occasionally defecated/urinated during odour extraction (Fig. 2e), further work is needed to determine the reliability of signalling in the A glomerulus for host discrimination.

## Robust decoding of human odour based on relative activity in H and U glomeruli

The narrow tuning of the H glomerulus to the long-chain aldehydes enriched in human odour generates a clear neural signal that *Ae. aegypti* mosquitoes may use for host discrimination. To gain more insight into how this signal can be used, either alone or in combination with signals from the U glomerulus, we trained a support-vector machine (SVM) linear classifier to identify samples as human or animal based on various glomerular inputs. When confronted with single-trial data from diverse odour samples delivered at the same 1X concentration (Fig. 6a, lighter points), a read-out of activity in H alone was enough to achieve over 90% accuracy (Fig. 6b top). However, when concentrations were allowed to vary (Fig. 6a, darker points), high doses of animal odour often evoked as much or more activity in the H glomerulus as lower doses of human odour, leading to mediocre accuracy based on H activity alone (75%; Fig. 6c top). In this case, a model in which activity in H is normalized by that in U provided clear improvement (83% accuracy; Fig. 6c middle) and was just as accurate as the less constrained model that took activity data from H and U as separate inputs (Fig. 6c bottom).

**Fig. 6.**
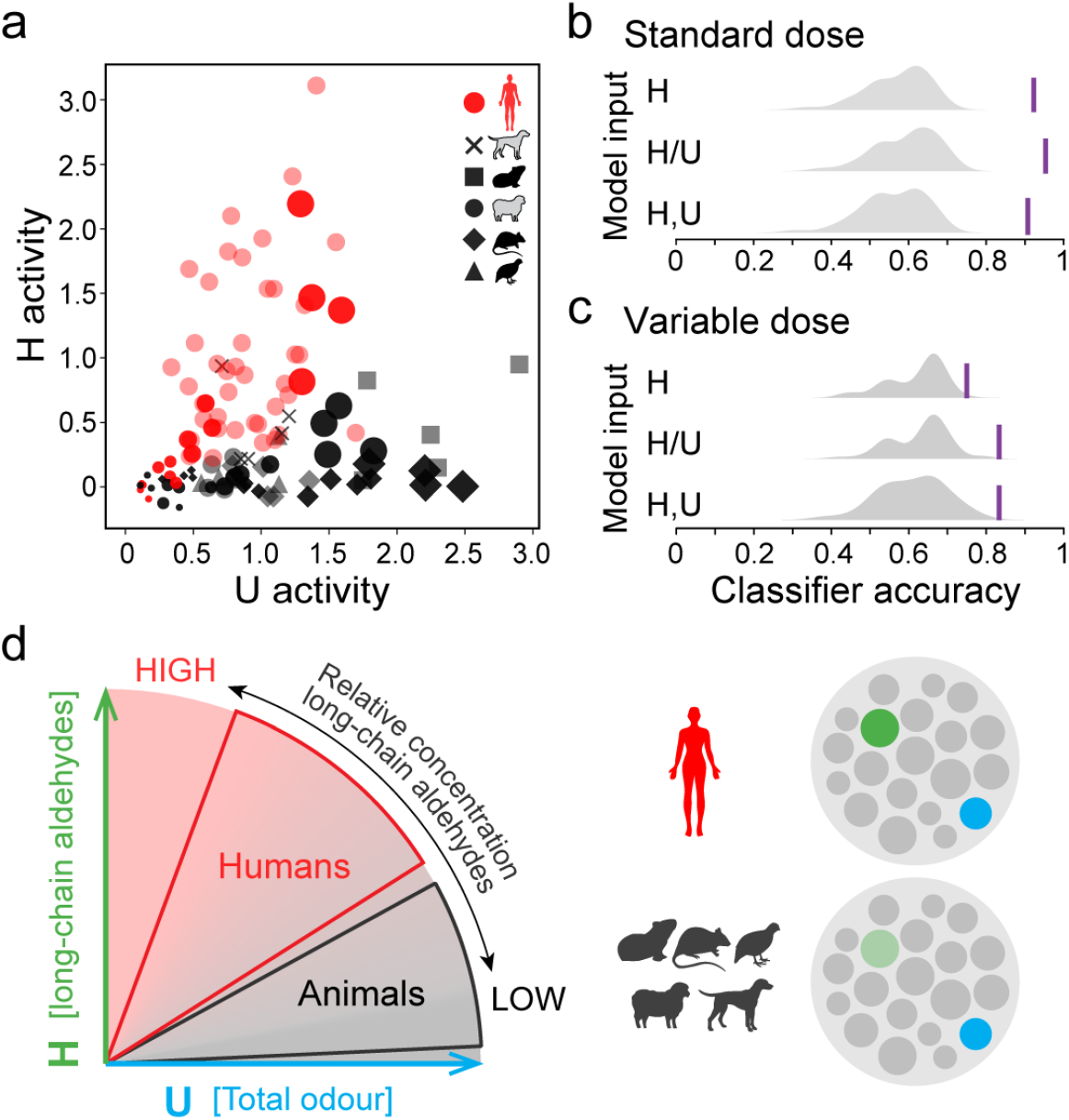
A robust neural code for human odour based on relative activity in H and U glomeruli. **a**, Single-trial data from the blend imaging experiments in Fig. 3. Lighter symbols, diverse human and animal blends delivered at 1X concentration (Fig. 3h–k); darker symbols, human, sheep, rat odour delivered at 4 concentrations (Fig. 3b–g). **b**, Accuracy with which an SVM linear classifier (see Methods) was able to discriminate between human and animal odour samples from the standardized-dose data (**a**, lighter symbols) using various configurations of glomerular input. Purple line, observed accuracy. Grey shading, distribution of accuracy values from 100 shuffled datasets. **c**, Same as (**b**) but for variable-dose data (**a**, darker symbols). **d**, Proposed model for human-odour coding in *Ae. aegypti*, wherein a comparison of activity in H and U glomeruli indicates the relative concentration of long-chain aldehydes in a complex blend and thus robustly differentiates human and animal odours.

These results suggest an interesting model for humanodour coding, wherein the broadly tuned U glomerulus provides a universal signal of total odour concentration, in comparison to which activity in H reveals the relative concentration of long-chain aldehydes in a complex blend (Fig. 6d). Only human odour, enriched in decanal and undecanal, evokes similar levels of activity in both glomeruli. As an indicator of relative concentration, this simple neural code is robust to variation in absolute odour levels and could allow mosquitoes to recognize humans in diverse contexts. The consistency of the difference in aldehyde content between human and animal blends also makes it robust to odour variation among individual humans and diverse animal species. Note that the U glomerulus is not the only signalling channel mosquitoes may use to achieve concentration-invariant coding. More selective channels with consistent responses to vertebrate odour (*e.g*. the palpal glomerulus that responds to 1-octen-3-ol or as-yet-unidentified IR+ glomeruli that respond to acids/amines) may also suffice.

## Discussion

Human-specialist mosquitoes use many different olfactory pathways (as well as thermal and visual cues) to find humans^72–74^. Carbon dioxide- and acid-sensing neurons are critical for baseline attraction^28,36,42–44^. However, the Orco+ neurons studied here are required for host discrimination^45^. The unique pattern of human-odour-evoked activity we document may thus be a dominant driver of the human-biting preference that makes *Ae. aegypti* such an efficient disease vector. Future work should explore ways to directly manipulate and test the effects of signalling in these glomeruli on behaviour. It will also be interesting to investigate integration of Orco-based signals with those from other olfactory pathways in the antennal lobe and higher brain areas. Finally, large-scale chemical screens for activators or inhibitors of H and/or U glomeruli may yield useful next-generation tools for mosquito control.

More broadly, our work provides insight into basic principles of olfactory coding. Previous studies demonstrate how olfactory systems may flexibly encode stimuli for learned associations^1–6^. In this case, broad activity across many glomeruli allows a large number of stimuli to be encoded by a limited set of coding channels. Simple stimuli with important innate meaning, in contrast, are often encoded by single, dedicated glomeruli. We know less about how animals evolve to encode complex blends they must recognize without prior experience. Like human and animal odours, many such stimuli are recognizable only by the specific combinations or ratios in which different blend components are mixed. At first glance, we might expect such complexity to also require complex combinatorial codes of differential activation across many glomeruli (*e.g*. Fig. 1f right). Instead, we see two prominent responses that could alone suffice – one accounting for overall odour concentration and a second narrowly tuned to shared compounds found at diagnostic relative concentrations (Fig. 6d, right). The unexpectedly sparse antennal lobe activity found in our study may be a general feature of innate olfactory responses.

Given the many compounds whose relative abundance distinguishes human and animal odours (Fig. 4e), why might *Ae. aegypti* rely heavily on aldehydes for discrimination? While other compounds almost certainly contribute^24,62,63^, aliphatic aldehydes are abundant in all vertebrate odours examined thus far and have been shown to affect host seeking in diverse mosquito taxa^26,75,76^. Even malaria parasites and orchids deploy them as mosquito attractors^77–80^. If aldehyde-sensing neurons are a conserved feature of a host-odour-attraction circuit in mosquitoes, re-tuning such neurons to the specific long-chain aldehydes enriched in human odour may have been one of the simplest evolutionary paths to human preference available to the ancestors of *Ae. aegypti*. Taken together, our work provides new insight into both mosquito preference for humans and the neural coding of complex olfactory stimuli that animal brains evolve to discriminate.

## Supplementary tables

**Supplementary Table 1.** Proportional abundance of all quantified compounds for 22 human samples (from 16 distinct individuals, see Methods), 5 animal samples, and 2 nectar-related samples.

**Supplementary Table 2.** All compounds quantified in the stimulus-odour analysis (Fig. 4), used for single-odorant imaging (Fig. 5, Extended Data Fig. 8), or used as reference odorants (Extended Data Fig. 2d). Identifying GC-MS characteristics of analytes and purities/vendors of synthetic compounds are listed where applicable. ‘Solvent’ refers to the solvent used for reference odorants and single odorant stimuli. ‘Dilution’ columns indicate the v/v ratio used for the initial panel of candidate reference odorants (L), the final panel of reference odorants (M), human odorants presented to mosquitoes at 1X human concentration singly or in combination (N and O, respectively, Fig. 5b–d, Extended Data Fig. 8a), and odorants presented at equal concentration (P, Fig. 5f–h, Extended Data Fig. 8b).

## Acknowledgements

We thank Vanessa Ruta, Leslie Vosshall, Mala Murthy, Emily Dennis, Jess Breda, Lu Yang, and members of the McBride lab for discussion and comments on the manuscript; Rickard Ignell for advice on odour extractions, odour analysis, and many related discussions; David Wevill of Markes for helping us adapt the thermal desorption system for stimulus delivery during neural imaging; Raphael Cohn and Alan Gelperin for advice on odour delivery systems; Silke Sachse, Ahmed Mohamed, Diego Pacheco, and David Deutsch for guidance on antennal lobe imaging; Hokto Kazama for advice on two-photon data analysis; Jonathan Pillow for advice on neural decoding analysis; Rob Harrell for mosquito embryo injections; Summer Kotb for help with human odour collections; and Howell Living History Farm, Nassau Park Pavilion PetSmart, and several dog owners for wool/hair samples. This work was funded in part by the National Institutes of Health via grants from NIDCD (R00DC012069) and NIAID (DP2AI144246) to C.S.M., and by the German Research Foundation (Deutsche Forschungsgemeinschaft) via a grant to D.M. (ME3737/3-1). C.S.M.’s laboratory is also supported by a Pew Scholar Award, a Searle Scholar Award, a Klingenstein-Simons Fellowship, a Rosalind Franklin New Investigator Award, and the New York Stem Cell Foundation. C.S.M. is a New York Stem Cell Foundation – Robertson Investigator.

## Author contributions

Z.Z. and C.S.M. conceived the project and designed and interpreted all experiments, with equal contribution from J.L.Z. on the odour analysis presented in Fig. 4. Z.Z. executed the experiments in Fig. 2, 3, 5, 6 and helped execute the experiments in Fig. 4; A.I. executed the experiments in Fig. 1a–d; B.J.M. and M.A.Y. provided advice on sgRNA and donor plasmid design for targeting the *orco* locus; A.L.K. helped design and execute the experiments in Fig. 4; J.L.Z. analysed the data presented in Fig. 4, Extended Data Fig. 6; S.T. designed and built the two-photon microscope used for volumetric imaging; M.S. developed the automated analysis pipeline for volumetric imaging, which he discussed with D.M.; Z.Z. and C.S.M. wrote the paper with help from J.L.Z. and other authors.

## Methods

### Ethics and regulatory information

The use of live non-human animals and non-human animal hair in olfactometer trials and odour extractions was approved and monitored by the Princeton University Institutional Animal Care and Use Committee (protocol #1999-17 for live guinea pigs and rats; #2113-17 for live quail; #2136F-19 for animal hair). The participation of human subjects in this research was approved by the Princeton University Institutional Review Board (protocol #8170 for olfactometer trials, #10173 for odour extractions). All human subjects gave their informed consent to participate in work carried out at Princeton University. Human-blood feeding conducted for mosquito colony maintenance did not meet the definition of human subjects research, as determined by the Princeton University IRB (Non Human-Subjects Research Determination #6870).

### Mosquito rearing and colony maintenance

All mosquitoes used in this research were reared at 26°C, 75% RH on a 14:10 light/dark cycle. Larvae were hatched in deoxygenated water and fed Tetramin Tropical Tablets (Pet Mountain, 16110M). Pupae were transferred to plastic-bucket or bugdorm cages, and adults were provided access to 10% sucrose solution *ad libitum*. Females were allowed to blood-feed on a human arm prior to egg collection. The Orlando (ORL) laboratory strain was used for both host-preference-behaviour testing and the generation of the *orco-T2A-QF2-QUAS-GCaMP6f* transgenic strain.

### Host preference assays

We tested the host preference of mated, non-blood-fed, 7–14 day old females that had been housed overnight with access to water only (no sucrose). We used a two-port olfactometer for choice and no-choice tests involving live hosts (Fig. 1a,c) or sleeves/hair (Fig. 1b,d) as previously described^24^. We first acclimated 75–100 female mosquitoes in the olfactometer for 5 min, then opened a sliding door and activated a fan to pull air through the two host chambers and expose mosquitoes to host odour. Mosquitoes were able to fly upwind, sample the host-odour streams, and choose to enter either host port. After 6 min, we counted the number of mosquitoes trapped in each host port. For two-choice trials with live hosts, one chamber contained a human hand and arm up to the elbow (belonging to one of six 22–43 year old individuals: 3 female, 3 male; 3 Caucasian, 2 East Asian, 1 South Asian). The human exhaled gently at the opening to the chamber once every 30 sec to provide a source of breath. The other chamber contained a guinea pig (*Cavia porcellus*; one of two 4–5 year old pigmented females), rat (*Rattus norvegicus domesticus*; one of two 2–6 month old Sprague-Dawley males), or button quail (*Coturnix coturnix*; one 2–3 year old female). For two-choice trials with animal hair, one chamber contained an arm-length section of a nylon stocking (L’eggs knee highs, black, 100% nylon) worn on a human arm for 24 hours and then stored at −20°C (same human subjects as in live-host trials). The other chamber contained a fist-sized wad of sheep wool (*Ovis aries*; from one female Romney sheep) or dog hair (*Canis lupus familiaris*; from one of four pet dogs – one Portuguese Water Dog, one Bichon, one Yorkie, one Old English Sheepdog). Sheep wool and dog hair was obtained from freshly shorn animals (from a sheep shearer, from a dog-grooming salon, or directly from dog owners) and stored at −20°C in sealed glass jars or odour-resistant nylon bags for up to 8 months before use. Both human-worn sleeves and animal wool/hair were supplemented with 1 sec on/1 sec off pulses of synthetic C¤2 (~1200 ppm). No-choice trials included the human or animal stimulus in one port with the second port left empty (air only).

We used a beta-binomial mixed generalized linear model (R^83^ package *glmmTMB*^84^) to model the probability of an individual mosquito choosing human *versus* each animal species in two-choice tests while accounting for overdispersion caused by trial-to-trial variation. Animal host species was included as a fixed factor, and date and individual human as random factors. We then extracted the model-estimated mean probability of choosing human with 95% confidence intervals (R package *emmeans*^85^) and converted this probability (p) to a preference index (PI = 2p-1) for data visualization. For no-choice trials, we used the same type of model to estimate the probability of responding to the given host, with host species included as a fixed factor and date as a random factor. The *R* function *cld* was used for pairwise comparison of least-square means.

### Generation of orco-T2A-QF2-QUAS-GCaMP6f strain

We used CRISPR-mediated homologous recombination (as described^47^) to knock in the QF2 transcription factor^86^ followed by the QUAS promoter (9 copies) and GCaMP6f^50^ coding sequence into the endogenous *orco* (AAEL005776) locus of the *Ae. aegypti* genome. We designed an sgRNA targeting the last exon of *orco* (GTCACCTACTTCATGGTGT<u>TGG</u>, PAM sequence underlined), generated template DNA by primer annealing with the NEBNext High-Fidelity polymerase (NEB, M0541S), and carried out *in vitro* transcription using the HiScribe T7 Kit (NEB, E2040S) by incubating at 37°C for 8 hours. We purified the transcription products using RNAse-free SPRI beads (Agencourt RNAclean XP, Beckman-Coulter A63987) and eluted them in Ambion nuclease-free water (Life Technologies, AM9937). We constructed the *T2A-QF2-9xQUAS-GCaMP6f-3XP3-dsRed* donor plasmid (Fig. 2a) using the InFusion HD Kit (Clontech, 638910). To preserve the *orco* coding sequence, the final 6 codons downstream of the cut site were included in the donor plasmid 5’ of the T2A, with synonymous codon substitutions incorporated to protect the sequence from Cas9 cleavage and minimize homology between the plasmid insert and the targeted locus. Homology arms (~1 kb) flanking the Cas9 cut site were amplified from ORL-strain genomic DNA via PCR. We found two divergent *orco* haplotypes segregating in ORL at similar frequency and therefore generated two versions of the donor plasmid with distinct homology arms. These two donors were mixed together for embryo injection (450 ng/μL each), along with sgRNA (80 ng/μL) and Cas9 protein (300 ng/μL; PNA Bio, CP01-200). A total of 1500 ORL embryos were injected at the Insect Transformation Facility at the University of Maryland Institute for BioScience & Biotechnology, yielding two independent transgenic lines with the construct inserted into the two distinct endogenous *orco* haplotypes. Insertion sites and sequences were verified by PCR and sequencing. The two lines showed indistinguishable patterns of GCaMP expression in the brain and peripheral organs, so we focused on the one corresponding to the major *orco* haplotype found in the AaegL5 reference genome^87^. This line was outcrossed to ORL for 8–9 generations. All experiments were carried out in heterozygotes, which displayed normal fitness and olfactory behaviours including strong attraction to/preference for human odour (data not shown).

#### Plasmids and primers

Plasmid backbone: psL1180, linearized with restriction enzymes NsiI-HF (New England Biolabs #R3127S) and AvrII (New England Biolabs #R0174S).

sgRNA template: forward 5’-GAAATTAATACGACTCACTATAGTCACCTACTTCATGGTGTGTTTTAGAGCTAGAA ATAGC-3’, reverse 5’-AAAAGCACCGACTCGGTGCCACTTTTTCAAGTTGATAACGGACTAGCCTTATTTTAACT TGCTATTTCTAGCTCTAAAAC-3’

Donor plasmid homology arms haplotype 1: left arm, forward 5’-CAGGCGGCCGCCATAGAGTTTCGCTTTTCC ACGCG-3’, reverse 5’-CCCTCTCCCGATCCATCCTTGAGTTGAACGAGAACCATGAAGTAGGTGACGACC-3’; right arm, forward 5’-TGTATCTTATCCTAGTGTTGGTGCAGTTGAAATAATTC-3’, reverse 5’-TATTAATAGGCCTAGAACTTACTTAAATCTGTGAAATCTCAGACC-3’.

Donor plasmid homology arms haplotype 2: left arm, forward 5’-CAGGCGGCCGCCATATTCAACGAGAGAAACGAAAGTT-3’, reverse 5’-CCCTCTCCCGATCCATCCTTGAGTTGAACGAGAACCATGAAGTAGGTGACGACC-3’; right arm, forward 5’-TGTATCTTATCCTAGTGTTGGTGCAGTTGAAATAATTC-3’, reverse 5’-TATTAATAGGCCT AGTCCACCTACGTATCATGACTAG-3’.

Sanger sequencing verification: forward 5’-AGCTGACCCT GTTGGCTTAC-3’, reverse 5’-CTTCAGCTTCAGGGCCTT-3’.

### Characterization of GCaMP expression in *orco-T2A-QF2-QUAS-GCaMP6f* strain

#### Brain

Brain immunostaining was carried out as previously described^48^. Heads of 7–10 day old mated mosquitoes were fixed in 4% paraformaldehyde (Electron Microscopy Sciences, 15713-S) for 3 hours at 4°C. Brains were dissected in PBS and blocked in normal goat serum (2%, Fisher Scientific, 005-000-121) for 2 days at 4°C. We then incubated brains in primary antibody solution for 2–3 days, followed by secondary antibody solution for another 2–3 days at 4°C. Brains were mounted in Vectashield (Vector, H-1000) with the anterior or posterior side facing the objective. Confocal stacks were taken with a 20X or 40X oil lens with an XY resolution of 1024X1024 and Z-step size of 1 μm. Primary antibodies: rabbit anti-GFP (1:10,000 dilution, ThermoFisher, A-11122) and mouse NC82 (1:50 dilution, DHSB, AB_2314866). Secondary antibodies: goat-anti-rabbit Alexa 488 (1:500 dilution, ThermoFisher, A27034SAMPLE), goat-anti-mouse CF680 (1:500 dilution, Biotium, 20065-1) and goat-anti-mouse Cy3 (1:500 dilution, Jackson ImmunoResearch, 115-165-062). We also dissected and stained the brains of 7–10 day old females whose sensory appendages had been removed with sharp forceps or micro-knives 5 days earlier.

#### Peripheral organs

We removed the antenna, maxillary palp, or proboscis of 7–10 day old female mosquitoes with sharp forceps. We then dipped them in pure ethanol for ~15 sec and mounted them on slides in pure glycerol for direct confocal imaging.

#### Reconstruction of glomeruli

We manually traced and reconstructed glomeruli (n=54) according to a previously published atlas^59^ using the TrakEM2 package^88^ in ImageJ^89^, except for 5 glomeruli that we could not reliably identify (CD1–4, PD7). Additionally, we found that a large portion of the anterior AL (antennal lobe), previously termed the Johnston’s organ centre^59^ (JOC), was innervated by Orco+ axons, consistent with recent work in *Anopheles gambiae*^51^. We therefore divided the JOC into 9 glomeruli based on NC82 neuropil staining, noting that the glomerular boundaries in the JOC were less clear than in other AL regions. In describing the orientation of the AL (*e.g*. Fig. 2b, Fig. 3), we also named the axes differently from the previous atlas to be more consistent with *Drosophila*^90^ and a recent atlas in *Ae. aegypti*^60^; the ventral/dorsal axis was switched with the anterior/posterior axis.

### Two-photon imaging

#### Microscope design

We designed a two-photon microscope that incorporates both resonant scanning^91^ and remote focusing^92,93^ to achieve rapid, volumetric, *in vivo* neural imaging. Remote focusing allows rapid switching of the imaging plane by moving a small, lightweight mirror located upstream in the imaging path. This alternative focusing method does not involve mechanical movements near the specimen, thereby avoiding specimen agitation and permitting axial scan speeds faster than those associated with traditional piezo-objective units. In diagnostic tests, transition times for switching between two planes were less than 6 ms. The combination of an 8 kHz resonant scanner and remote focusing resulted in volumetric-stack-imaging speeds of 512 pixels x 512 lines x 10 planes at 3 Hz.

The microscope uses a pulsed (80 MHz) Ti:Sapphire laser (Coherent Chameleon Vision II) tuned to 920 nm, with laser intensity rapidly controlled on the μs timescale with a pockel cell (Conoptics 350-80-LA-02 KD*P). The beam entering the microscope is first sent through a half-wave plate (Thorlabs AHWP10M-980), a polarizing beam splitter cube (Thorlabs PBS252), and a quarter-wave plate (Thorlabs AQWP10M-980) before entering the remote objective (Olympus UPLFLN40X). It is then reflected on a 7 mm mirror (Thorlabs PF03-03-P01) mounted on a voice coil (Equipment Solutions LFA2004). The beam then crosses the remote objective and the quarter-wave plate in reverse direction before being reflected by the polarizing beam splitter cube. It then enters a non-magnifying relay telescope made of two identical achromatic lenses (Thorlabs AC254-150-B) that brings it to the scanning unit located in a plane conjugated to the remote focus objective back aperture. The scanning unit includes an 8 kHz resonant scanner (Cambridge Technologies CRS8) for the fast axis and a 6 mm galvanometer scanner (Cambridge Technologies 6215H). The beam then travels through the 150 mm scan lens (Thorlabs AC508-150-B) and a 200 mm tube lens (2 identical lenses, Thorlabs AC508-400-B) to reach the imaging objective (Olympus LUMPLFL 40X Water, NA 0.8), whose back aperture is conjugated to the scanning unit. The distance between the scan lens and the tube lens is precisely set to be the sum of their respective focal lengths, a condition that minimizes optical aberrations when using remote focusing^92,93^. The microscope’s field of view is 550 μm in diameter.

The quantity of glass present in the optical path of this microscope generates significant group-delay dispersion, for which the laser internal pre-compensator cannot fully compensate. This results in lower fluorescence excitation. We therefore added another compensator made of a pair of SF10 prisms (Newport 06SF10), through which the beam passes before entering the microscope. We adjusted the distance between prisms to roughly maximize the fluorescence signal, then relied on the laser internal precompensation unit for fine maximization.

The fluorescence signal is separated from the laser path by a dichroic mirror (Semrock FF670-SDiO1) and detected by GaAsP photomultipliers (PMT; Hamamatsu H10770PA-40) after successively passing through a multiphoton short-pass emission filter (Semrock FF01-720sp), a dichroic mirror (Semrock FF555-Dio3), and a band-pass filter (Semrock FF02-525/40-25 for the green channel; Semrock FF01-593/40-25 for the red channel). The PMT output signals are amplified (Edmund Optics 59-179) and digitized (National Instrument PXIe-7961R FlexRIO). The microscope is controlled by the ScanImage (Vidrio) software using additional analogue output units (PXIe-6341, National Instruments) for the laser-power control, the scanners control, and the voice-coil control.

#### Mosquito preparation

We custom-designed a mosquito holder with a 3D-printed plastic frame and thin stainless-steel plate (Fig. 2c, thickness 0.001 inch, design files available at https://github.com/mcbridelab/Zhao_2020_HumanOdorRepresentation). A tiny mosquito-head-sized hole was photo-chemically etched on the plate (ETCHIT Company). To prepare for imaging, we anaesthetized a female on ice for ~1 min, pushed the anterodorsal side of her head into the hole, and fixed it with UV glue (RapidFix 6121830ES). The antennae, maxillary palps, and proboscis remained below the metal plate and contacted neither the plate nor the glue. We added room-temperature saline (103 mM NaCl, 3 mM KCl, 5 mM TES, 26 mM NaHCO_3_, 1 mM NaH_2_PO_4_, 1.5 mM CaCl_2_, 4 mM MgCl_2_, 10 mM Trehalose, 10 mM Glucose; pH 7.1) to the holder and used sharp forceps to remove a section of the head capsule (including both cuticle and the edge of the eyes) from the part of the head protruding through the metal plate. During imaging, we continuously perfused saline bubbled with carbogen (5% CO_2_, 95% O_2_) through the holder and across the open head capsule at 125 mL/hour.

#### Data acquisition

We used the *ScanImage*^94^ package in Matlab to control the microscope and acquire images. For each individual, we chose either the right or left antennal lobe and recorded movies of odour-evoked activity (starting 7–30 sec before and continuing 20–60 sec after syntheticodorant puffs; ~30 sec before, ~140 sec after puffs of complex extract) that covered the entire structure in 22 stacks, 4 μm apart, at 128 x 128 pixel resolution. The resulting voxel size was approximately 0.9 x 0.8 x 4 μm^3^, and the volumetric imaging rate was 3.76 Hz. We increased the laser power exponentially with depth (ranging from 7.5 to 10 mW) to account for light decay and scattering in deeper tissues. Laser power as measured at the sample plane was 10 mW. After recording odour-evoked activity, we acquired 30–40 high-resolution structural volumes at high laser power to aid registration and downstream analysis. For this, we imaged the AL in 120-180 z-stacks, 1 μm apart, at 256 x 256 pixel resolution.

### Two-photon data analysis

#### Reference-odorant selection

We selected reference odorants based on a preliminary imaging data set comprising the glomerular responses to 60 candidate odorants (n=2 mosquitoes, Supplementary Table 2). After extracting glomerular signals, we obtained a (glomerulus x odorant) matrix A of mean odorant responses. We then employed the ConvexCone algorithm (see below) to select c=1,…,N columns (corresponding to odorants) from A into a series of matrices C_1_,…, C_N_ and measured the respective norm error ||A - C_c_ X_c_||. This norm error decreased quickly with increasing c (Extended Data Fig. 2d), suggesting that a small subset of odorants can account for a large part of the matrix norm (variance/glomerular activity). Accordingly, we chose the 11 odorants that best reduced the norm error, along with three odorants that were of special interest (sulcatone, 1-octen-3-ol, phenylethylamine), as reference odorants for all subsequent imaging (Supplementary Table 2).

#### Motion correction and morphological registration

We first performed 3D motion correction on each volumetric movie of odour-evoked activity using the NoRMCorre package^95^. We then used the *warp* function in the Computational Morphometry Toolkit (CMTK, http://nitrc.org/projects/cmtk) to correct for potential motion and brain deformation between movies from the same brain. We created the two-photon AL template by iteratively registering and averaging the ALs from 13 high-quality brains with the CMTK *warp* and *avg_adm* functions. We registered each AL to the two-photon template, again using the CMTK *warp* function, so all brains were aligned in the same coordinates and had similar shape (Extended Data Fig. 2a).

#### Unsupervised segmentation based on activity

An odourresponse recording R_i_ contains a 3D volume for each time point: it is a tensor with three spatial dimensions (x=128, y=128, z=24) and one time dimension: R_i_=(1:x, 1:y, 1:z, 1:t). For each time point t_p_, we performed spatial smoothing of R_i_(:,:,:, t_p_) with a 3D Gaussian kernel. We used a movingaverage filter for temporal smoothing along the time dimension of R_i_. A mask M covering the AL served to cut out the background by element-wise multiplication with each R_i_. Due to elevated baseline calcium levels within the AL area, we could obtain a mask by simple Otsu thresholding of an average volume for the pre-stimulus interval. For each AL, we extracted functional clusters, *i.e*. clusters of voxels with correlated activity in R = [R_1_,…, R_N_odours_]. These clusters can be interpreted as glomeruli, especially if they have a spatial-functional match in another AL. For functional clustering, we employed a non-negative matrix-decomposition scheme solved with the ConvexCone algorithm^96^ that has a track record of successful application to imaging data from different species. Briefly, R is reshaped to a matrix A with m = x * y * z rows (voxel vectors) and n = odours*time columns (time series vectors). We then decompose A into a matrix of the c most relevant time series C ∈ A and their spatial mappings in X, such that || A - CX||_Fr_ is minimized subject to a non-negativity constraint on X. In practice, this is carried out on a rank k=50 representation of A obtained by SVD.

The continuous-valued and non-negative X acts as a fuzzy cluster membership indicator, locating the time series signals from C in space and also encoding cluster overlap due to signal mixtures, *i.e*. a voxel can ‘belong’ to several clusters to different degrees. For creating the 3D AL map visualizations (Extended Data Fig. 2), we binarized the continuous X with algorithm 2 from a previous study^97^. The solid clusters obtained were then rendered as 3D objects. We observed cases of apparent overclustering, where two or more overlapping clusters with distinguishable, but nevertheless similar, odour responses were found. This could be due to the ability to resolve actual glomerular subcompartments, or due to signal bleed-through from upper layers, which creates different signal mixtures in different subvolumes of a glomerulus. In order to resolve overclustering, we merged clusters that overlapped in the 3D AL maps and had odour-response profile (mean df/f responses) correlations greater than 0.7.

#### Automated glomerulus matching across brains

We matched glomeruli across brains if they were similar in both odour-response properties and relative position, allowing for a certain degree of physiological and anatomical variation (Extended Data Fig. 2c, Extended Data Fig. 5b,c). To simplify functional and spatial comparisons, we compressed the glomerular time series to the mean df/f responses for each odour. We then performed pairwise matching of all subject ALs to a single target AL. Matching a subject to the target can be cast as an assignment problem on a bipartite graph G = (V=(S, T), E), where the glomeruli in the subject and target AL are represented as vertices S and T, respectively, that have to be connected by a set of edges E in a way that optimizes a cost criterion. For a given cost criterion, the optimal assignment can be computed with the Hungarian algorithm^98,99^. We minimized the (weighted) functional-spatial distance d(a,b) = w_f_ * d_functional_(a, b) + W_s_ * d_spatial_(a, b), where d_functional_(a,b) denotes the Euclidean distance between the odour-response profiles (mean df/f responses) of glomeruli a and b, while d_spatial_(a, b) refers to the Euclidean distance between the centroids of glomeruli a and b that have been pre-registered into the same 3D space. All distances were normalized to remove scaling differences between functional and spatial distances.

Global optimization of d leads to a complete assignment of all glomeruli from S to all glomeruli from T. However, due to missing (glomeruli that were not detected) or additional clusters (overclustering or non-glomerular clusters) in either brain, not all glomeruli may have a meaningful match. We thus employed the Hungarian algorithm to compute all glomerulus matches that are feasible under functional and spatial constraints:

- d is set to infinity if d_spatial_(a, b) > c_spatial_ (*e.g*. the diameter of a typical glomerulus)
- d is set to infinity if the odour response profile correlation corr(a, b) > c_functional_ These constraints specify the criteria we demand for an acceptable match (in terms of response similarity and spatial distance), while the optimal assignment under these constraints is left to the algorithm. Whenever the constraints led to infeasible matches, the respective subject AL glomeruli were excluded from further analysis.

#### Across-matrix PCA of common odour response space

We also pursued the alternative strategy of constructing a common odour-response space for all mosquito brains, allowing us to visualize distances between odour responses in a way that is unaffected by parameter settings (such as the number of clusters) or possible matching errors of the glomerulus-matching approach (Extended Data Fig. 5a). There are N_odours_ odour-response recordings for the j*th* mosquito brain, R^(j)^=[R_1_,…, R_N_odours_], where the sequence of odours is assumed to be the same across brains. After preprocessing the R^(j)^ (see above), we performed df/f normalization for all voxel time series in the R^(j)^, followed by spatial median filtering to remove a few extremely high df/f values in individual voxels. The time series in the R^(j)^ were then reduced to mean odour responses during an interval after odour presentation and reshaped to matrices A^(j)^ with m=#voxels rows and n=#odours columns. We then constructed A^(all)^ = [A^(1)T^ …, A^(N_brains)T^]^T^, *i.e*. the (N_voxels_ * N_brains_) x N_odours_ matrix containing the row-concatenated and whitened matrices A^(j)^. The principal components of A^(all)^ are across-matrix principal components that span a common odour-response space for all brains. PCA provides the best rank-k approximation to A^(all)^ in the sense that it finds matrices U,V that minimize || A^(all)^ - UV |_Fr_, where V is a k x (N_voxels_* N_brains_) matrix that can be partitioned as V=[V^(1)^,…, V^(N_brains)^]. By projecting the A^(j)^ onto the V^(j)^, we can obtain the positions of the odour/brain combinations in the space of the top-k across-matrix principal components.

#### Manual identification of target glomeruli

The three target glomeruli could be reliably identified across brains based on position and responses to key reference odorants (Extended Data Fig. 5c, Supplementary Table 2). Human-sensitive glomerulus H was located in the anterior AL, adjacent to a landmark Orco-glomerulus in our two-photon images (a black hole surrounded by Orco+ glomeruli; Fig. 2b; Extended Data Fig. 2a-b), and responded to 10^-2^ heptanal. Animal-sensitive glomerulus A was located in the dorsal AL and responded to 10^-2^ phenol. Universal glomerulus U was located in the posterior-medial AL and responded to 10^-2^ benzaldehyde. Glomeruli H, A, and U tentatively correspond to V3, MD2, and PD1, respectively, in a previously published atlas^59^, but we cannot be sure without molecular markers.

#### Summarizing neural activity

We used area under df/f curves as a metric for neural activity. For single-odorant stimuli, which evoked single df/f peaks, we defined the peak boundaries by first locating the max (for activation) or min (for inhibition) points in the df/f curve and then extending from the max/min points until df/f dropped to background levels. For odour extracts, which sometimes evoked multiple peaks, we integrated df/f values from desorption of the focusing trap to the end of neural recording (0 to 140 sec). To account for variation in responsiveness across brains, we normalized area values for single odorants by the response of glomerulus H to decanal (at whatever concentration was used in that experiment) and for odour extracts by the strongest response evoked in a given brain by any odour extract used in the experiment (min-max normalization, where min is zero). We used the paraView software to render the neural responses into 3D plots^100^.

#### Neural decoding analysis

We trained support-vector machine (SVM) classifiers to test whether responses to human odour could be reliably discriminated from responses to animal odour on a trial-by-trial basis. We used activity or relative activity of selected glomeruli as the input variables, along with mosquito brain identity as an indicator variable. Models were constructed and trained with the *SVC* function in the *scikit*-learn Python module^101^. We evaluated the accuracy of classifiers using the leave-one-out approach. Data were randomly shuffled 100 times to obtain the null distribution of classifier accuracy.

### Headspace odour extraction

#### Extraction from human volunteers

We modified a previously published protocol for human-body headspace odour extraction^102^. Subjects were asked to bathe using an unscented soap (365 Everyday Value 0-99482-41321-7) 3 days before odour collection and then avoid the use of all soaps, skin products, swimming pools, and hot tubs thereafter. Subjects were also asked to avoid all water baths/showers, spicy food, and alcohol for 24 hours before collection. At the time of extraction, subjects lay nude inside a custom-made 80” x 48” Teflon FEP bag with 4 ports on each side (Fig. 2e middle, American Durafilm) and covered by a privacy blanket over the bag. The bag’s opening was loosely cinched around each subject’s neck, and a pair of Tenax TA tubes (Markes International Inc., C1-CAXX-5003) was inserted into each of the 8 ports. A Teflon tube pushed into the bag through the neck hole provided a source of charcoal-filtered zero-grade air at 3.6 L/min (filter: Whatman 67221001; air: Airgas AIZ300). Two vacuum pumps (KNF Neuberger, UN811KV.45P115V) were used to pull air out of the bag through the 8 ports at 400 mL/min per port (200 mL/min per Tenax tube) for 2 hours while the subject watched a movie or listened to music. We confirmed that the Tenax tubes captured all major odorants (no breakthrough) at the given flow rate and duration in test extractions where two tubes were placed in series. The bag was washed (Babyganics fragrance-free dish soap and DI water) and autoclaved before each use. Three subjects underwent replicate extractions weeks to months apart (B37, L42, Q82; see Supplementary Table 1), demonstrating moderate within-individual consistency of the odour profile over time (Extended Data. Fig. 6e).

#### Extraction from non-human animals, plants, and honey

We collected headspace odour from quail, guinea pigs, rats, and milkweed flowers using custom-designed glass extraction chambers (Fig. 2e top; 14 cm diameter x 24 cm long or 8 cm diameter x 30 cm long). A single rat, a single guinea pig, a pair of quail, or several freshly cut milkweed inflorescences (*Asclepias syriaca*, leaves removed) were placed in the chamber for each extraction. We collected headspace odour from sheep wool, dog hair, and honey using a 250 mL glass gas-washing bottle (Fig. 2e bottom, Chemglass, CG-1112-02). Twenty-five grams of hair were packed into the bottle and heated to approximate homeothermic-vertebrate body temperature in a 37°C water bath. For honey, we smeared 25 grams (Tasmanian Honey Company, leatherwood honey) across the inner face of the bottle without heating. We used the same live animals and hair described under ‘Host preference assays’ except that there were two quail instead of one and the pet dogs were a German Shepherd, a Yorkie, and two Portuguese Water Dogs. All extractions were supplied by charcoal-filtered tank air that was pulled through one or two Tenax TA tubes (Markes International Inc., C1-CAXX-5003) using a vacuum pump (Sensidyne, Gilian 800i). Flow rate and extraction duration: rats, 200 mL/min through 1 Tenax tube for 2 hours; quail, 200 mL/min through each of 2 Tenax tubes for 2 hours; guinea pigs, milkweed, sheep wool, dog hair, and honey, 400 mL/min through 1 Tenax tube for 1 hour. We confirmed that the Tenax tubes captured all major odorants (no breakthrough) at the given flow rate and duration in test extractions where two tubes were placed in series. The glass extraction chamber and gas-washing bottle were washed (Babyganics fragrance-free dish soap and DI water) and rinsed with methanol (HPLC-grade, ≥99.9%, Sigma Aldrich) and hexane (≥99.8% for GC-MS, SupraSolv) before each extraction.

Animal waste was sometimes present in the odour-extraction chambers for guinea pig, rat, and quail and thus may have contributed to the corresponding odour samples and to the list of animal-enriched compounds^15,71^ (*e.g*. p-cresol in Fig. 4e). However, sheep odour, which came from unsoiled wool, was nested among the other animals in almost all compound-specific analyses (Extended Data Fig. 6f).

#### Processing of odour extracts

We generated 16 Tenax tubes for each individual human (Fig. 2e middle). Since tubes had the potential to vary depending on their position in the extraction bag, we decided to pool the 16 tubes into 4 more-homogeneous aliquots using the ‘stacking’ feature of our Markes thermal desorption system (see Delivery of host-odour extracts via thermal desorption, below; Extended Data Fig. 3e). Each of the 4 aliquots represented a pool of one tube from each of the four bag positions (shoulder, waist, knee, foot). We then used the Markes system to puff a small portion of the first aliquot to a new Tenax tube for GC-MS analysis and reserved the remaining samples (1 partial aliquot + 3 full aliquots) for imaging. We similarly stacked and aliquoted the 12-16 tubes obtained for each animal species and for honey. For these samples, each tube or pair of tubes came from a separate extraction and had the potential to vary due to individual or day-to-day variation. For milkweed, we used a single tube without stacking due to the high odour concentration.

### Odour analysis

#### TD-GC-MS analysis

We used an Agilent GC-MS system (Agilent Technologies, GC 7890B, MS 5977B, high-efficiency source) outfitted with a DB-624 fused-silica capillary column (30 m long x 0.25 mm I.D., d.f.=1.40 μm, Agilent 122-1334UI). Tenax tubes were inserted into a Gerstel TD3.5+ thermal-desorption unit (Gerstel Inc.) mounted on a PTV inlet (Gerstel CIS 4) with a glass-wool-packed liner. Tubes were heated in the TD unit from 50°C to 280°C at a rate of 400°C/min, then held at 280°C for 3 min. During the TD heating time, volatiles were swept splitless into the cold inlet (−120°C) under helium flow of 50 mL/min. After the tube was removed and the inlet repressurized, the inlet began heating at a rate of 720°C/min to a 3 min hold temperature of 275°C. The GC oven program began simultaneously with inlet heating, starting at an initial temperature of 40°C and ramping at a rate of 8°C/min to a 10 min hold temperature of 220°C. Transfer from the inlet to the GC column was performed at a 20:1 split ratio (40:1 split for milkweed). Carrier-gas flow rate was 40 cm/s. The MS was operated in EI mode, scanning from m/z 40 to 250 at a rate of 6.4 Hz.

#### Peak detection

The major steps in our analysis pipeline are illustrated in Extended Data Fig. 6a. We imported the raw GC-MS data into Agilent’s MassHunter Unknowns Analysis program (version B.09.00, build 9.0.647.0) and extracted peaks using the built-in deconvolution algorithm with window sizes of 25, 50, 100, and 200. We also extracted peaks using the TIC (total ion chromatogram) analysis option with a window size of 100. The deconvolution algorithm looks for correlated peaks in ion abundances and can pull apart partially co-eluting peaks. TIC analysis integrates the entire area under peaks in the total ion chromatogram. We used both algorithms in order to cast an initial broad net that would capture all potential peaks. In most cases of non-overlapping peaks, the deconvolution algorithm gave similar peak areas to TIC integration. Therefore, for the final dataset, we selected peaks found by TIC integration only in the very small number of cases where the deconvolution algorithm had clearly truncated the edges of the peak.

#### Preliminary compound identification

We identified peaks by using Unknowns Analysis to search the NIST17 MS EI library for matches. The program finds the best match in the reference library for each peak (with a minimum match score of 70), then for each compound selects the peak with the highest match score. We manually selected alternate best-hit peaks (sometimes with match score below 70) if the automated choice looked non-Gaussian or was composed of misaligned component-ion peaks (implying the peak was made up of multiple co-eluting compounds). We also manually selected alternate peaks if there was an excess of background ions in the automated choice. We ensured that retention times for each compound matched across samples.

#### Targeted search for 1-octen-3-ol

Because 1-octen-3-ol is known to be an important mosquito attractant despite its low abundance in host odours^68,69^, we performed a targeted search for this compound. In 19 out of 21 samples where 1-octen-3-ol was present, the level was too low to be detected by the automated pipeline, particularly because 1-octen-3-ol co-elutes with the common compound benzaldehyde. Therefore, we wrote custom R scripts to quantify the amount of 1-octen-3-ol in our samples. We used the *mzR* package^103^ to access the raw GC-MS data from mzxml files. For each sample, we fit Gaussian curves to the m/z 51 and m/z 57 ion peaks under the shared 1-octen-3-ol/benzaldehyde peak; at this retention time, these ions are diagnostic for benzaldehyde and 1-octen-3-ol, respectively. The abundance ratio of the two ions is directly proportional to the abundance ratio of the two compounds. We used this ratio to infer the amount of 1-octen-3-ol present in every sample by extrapolating from the two samples where the deconvolution algorithm successfully pulled out and integrated a separate 1-octen-3-ol peak.

#### Criteria for including compounds in the analysis

In the final dataset for analysis, we retained only compounds eluting between 6 and 22 minutes. We ignored early-eluting compounds because breakthrough analysis suggested we were not able to obtain quantitative estimates of their abundance. Few compounds eluted after 22 minutes; these tended to be less volatile and more difficult to identify because of the high background. We also removed obvious contaminants: siloxane column artefacts, 4-cyanocyclohexene (a compound likely from nitrile gloves^104^), and other components not plausibly produced by biological metabolism because they contained heteroatoms other than O, N, and S. Finally, we ignored carboxylic acids because (1) they are difficult to quantify reliably without derivatizing samples, and (2) they are not typically detected by the Orco+ neurons^105^ on which we focus in this study.

We set an abundance criterion for including compounds in the analysis: for a focal compound X to qualify, it must have constituted at least 2% of the ‘odour profile’ of at least one sample, where the ‘odour profile’ comprises noncontaminant compounds just as or more abundant than X in the given sample (Extended Data Fig. 6a). Because we had a large number of human samples, and in order to avoid unfairly including an excess of compounds prevalent among humans, we randomly selected a single sample (L95, see Supplementary Table 1) to serve as the human representative for this compound-qualification step. Forty-eight compounds met our criteria (Extended Data Fig. 6b,c) and were thus quantified across <u>all</u> samples, regardless of abundance in any given sample.

#### Verification with synthetic standards

We used retentiontime and mass-spectrum information from external standards to verify the identities of all compounds mentioned by name in this paper, except those marked by a parenthetical question mark in Fig. 4 and Extended Data Fig. 6f. Sources of external standards are listed in Supplementary Table 2. Of the 48 compounds included in the analysis, 25 were verified by external standards, 18 were identified based only on a mass-spectrum match to the NIST 17 library, 4 were assigned a compound class but not a precise identification based on mass-spectrum characteristics, and 1 remained unidentified. Even when the identity of a compound was uncertain, we were able to use retention time and mass spectrum to reliably locate it across samples.

### Odour stimulus preparation and delivery

#### Delivery of host-odour extracts via thermal desorption

We adapted a thermal-desorption (TD) system marketed for GC-MS applications to deliver complex odour extracts from Tenax tubes to mosquitoes during imaging. The Unity-Ultra-xr TD system from Markes International Inc. uses a 2-step desorption procedure (Fig. 2f). The sorbent tube containing the sample is heated slowly to a high temperature to desorb odorants, which are carried by nitrogen flow to a cold, sorbent-filled focusing trap. The focusing trap is extremely narrow and can therefore be heated ballistically (to 220°C in 3 sec) to release all odorants during a very short time window – more or less simultaneously. For GC-MS applications, the odorants then enter the GC, and the focusing step serves to narrow the GC-MS peaks. In our case, we connected the output flow of the TD system to the mixing manifold of our odour-delivery system (see below, *Delivery of synthetic odorants and blends*) and used a thermocouple thermometer (AMPROBE, TMD-52) to confirm that the final mixed flow (TD output + carrier air) was at room temperature. We then optimized puff shape and duration for imaging by increasing the flow rate through the cold focusing trap (to 30–120 mL/min depending on split-flow ratio) and setting a high dilution ratio at the mixing manifold (1:30, TD output to carrier air). Using a photoionization detector (PID; Aurora, 200B), we observed a single ~3 sec peak for single odorants, mixtures of two odorants, and human extracts (Extended Data Fig. 3a–c). However, not all compounds in the odour extracts are detected well by a PID. We therefore also collected on a Tenax tube the odour stream coming from the delivery system for 10, 20, or 30 seconds following the odour-release command and analysed the collected volatiles via GC-MS. Most components were released within the first 3–7 seconds (10 sec relative to the release command given a 3 sec delivery delay), with one or two late-eluting compounds requiring up to 17 seconds (20 sec with delay) to fully desorb (Extended Data Fig. 3d).

We relied on two additional features of the Unity-Ultra-xr TD system to precisely control and standardize odour stimuli across individual puffs and mosquitoes. First, we used the ‘stacking’ feature to pool multiple odour tubes from the same or different extractions and thereby generate concentrated, homogeneous extracts for each animal species or human individual (Extended Data Fig 3e; see also *Headspace odour extraction – Processing of odour extracts*, above). Stacking is achieved by desorbing tubes onto the focusing trap one after the other, while maintaining the trap at a constant, low temperature (30°C). The accumulated odour is then released via ballistic heating and collected on a new tube.

Second, we used the ‘split-recollect’ feature to dispense concentration-matched aliquots for use in imaging (Fig. 2f–g; Extended Data Fig. 3e,f). This feature allowed us to puff a specific percentage of a sample to a new tube (or the mosquito) by splitting the flow during desorption of the focusing trap. Moreover, the leftover portion can be recollected for future use. The percentage puffed is precisely determined by the split-flow ratio. We adjusted the ratio based on the total volatile content of a stacked sample (estimated via prior GC-MS analysis) to generate aliquots of equivalent dose across diverse odour stimuli (Fig. 2g; Extended Data Fig. 3e).

We also used the split-recollect feature to deliver a prespecified percentage of each concentration-matched aliquot to the mosquito during imaging and recollect the remainder. We desorbed the sample tube for 2 min with the temperature ramping to 200°C and then desorbed the focusing trap for 1 min at 220°C (Extended Data Fig. 3d). The trap was then cooled to 30°C over a period of 25 sec, with continuing nitrogen flow, before the output valve closed. We adjusted the split-flow rate (fraction of odour puffed *versus* recollected) to standardize dose across mosquitoes. For each mosquito, we randomized the order of delivery of stimuli and waited for approximately 10 min between stimuli. We used a PID and GC-MS to verify that both the absolute quantity of odour extract delivered and the ratios of constituent components were stable across at least 10 replicate puffs using this split-recollect feature (Fig. 2h; Extended Data Fig. 3f).

To define a standard dose, we selected one reference subject whose total odour content was approximately average among all human subjects. We then defined 1X human dose such that the release rate from the odour-delivery system was approximately equal to the release rate from the reference subject’s whole-body odour extraction. This calculation took into account the duration of the odour extraction, the number of collection tubes, the duration of the odour puff, and dilution by the carrier stream in our odour-delivery system. 1X doses of other stimuli were defined as having the same total odour content as 1X reference human.

#### Selection of synthetic odorant panel

We focused on commercially available major compounds (>3%) in any species, ignoring compounds that met this threshold in only one or two humans. Some minor compounds above this threshold were not included because they were under or close to the 3% cut-off in the initial odour analysis available at the time we were choosing individual odorants for imaging. None of these compounds was above 5% in any sample in the final analysis that quantified 48 compounds.

We excluded geranylacetone because it was unstable in solution and 1-dodecanol because it was not found in our initial dog sample (a single individual instead of an average over 4 dogs). We also included pentanal and undecanal despite their low abundance in order to test the full complement of straight-chain, saturated aldehydes found in our samples. Altogether, our panel comprised 19 compounds. These compounds composed 72–99% of the quantified odour content of our samples (with the exception of honey, 45%).

#### Estimation of vapour-phase concentration of synthetic odorants and blends

We developed a method to measure the volatility of single odorants based on a previously published study^70^ (Fig. 5a). We first made standard dilutions of single odorants in paraffin oil or water. For a given odorant, the exact dilution ratio (neat, 10^-2^, 10^-3^, or 10^-4^) was chosen based on published or predicted vapour pressure (SRC PHYSPROP Database) in order to keep concentrations within a similar range appropriate for GC-MS analysis. We then used our high-throughput odour-delivery system (see next section) to sequentially puff the odorants to a conditioned Tenax tube (instead of to a mosquito). Each Tenax tube contained single puffs of 4–6 different odorants, and each odorant was puffed to 3 independent tubes. We then analysed the tubes via TD-GC-MS, quantified the peak area for each odorant, averaged across replicates, and back-calculated the volatility of each odorant in our set-up.

#### Delivery of synthetic odorants and blends

We designed and built a high-throughput system to deliver synthetic odorants and blends during imaging. Our design was inspired by the commercial Aurora 206A system, but has 20 odour channels and a flush stream. We briefly describe the system here, with more detail in Extended Data Fig. 7. The system includes a humidified carrier air stream, an odour stream with separate channels for twenty 40 mL odour-dilution bottles (Scientific, 12-100-108), and a CO_2_ stream. The odour and CO_2_ streams are also each coupled to their own control stream that serves to equalize total flow when the stimulus is not being delivered. A final high-flow flush stream purges the flow path of the odour stream between puffs to remove traces of the previous stimulus. Mass-flow controllers (Aalborg, GFCS-010007 and GFCS-010008) dynamically regulate the flow through all streams except the flush, and a PTFE manifold (Cole-Palmer, EW-31521-13) acts as a final mixing station. All valves (3-way, Cole-Palmer UX-01540-11; 2-way, Pneumadyne S10MM-20-12-3 and MSV10-12) and mass-flow controllers are controlled by Arduino boards (Arduino Mega 2560 r3, Uno r3). We wrote an open-source GUI in Python to control the odour-delivery system and trigger image acquisition (https://github.com/mcbridelab/Zhao_2020_HumanOdorRepresentation). We prepared an odorant panel by filling each of the twenty 40 mL odour vials with 3 mL of odorant dilution. When switching in a new odorant panel, we flushed the flow path of the system with hexane and purged it overnight with filtered air to remove potential traces of the previous panel. We characterized the puff shape (Extended Data Fig. 7b) and long-term stability (Extended Data Fig. 7c) of our system using a PID, with 2-heptanone as the test odorant.

During imaging, we recorded the neural response of each mosquito to 2–3 replicate, 3-second puffs per odorant, presented in random order with an inter-puff interval of 60–90 sec. We also recorded the baseline response for a given odour channel (clean air passing through the channel’s valves/tubing but bypassing the odour bottle) and the response to solvent only and subtracted these from the response to stimulus. We used odorants of the highest commercially available purity and diluted them in paraffin oil (Hampton Research, HR3-421) or ultrapure water (Supplementary Table 2).

## Data availability

All relevant data supporting the findings of this study are available from the corresponding authors upon request. For odour-profile analysis, data are included with the paper as Supplementary Table 1.

## Code and reagent availability

Code used for analyses and all unique biological materials generated in this study are available from the corresponding authors upon request. Control code for the odour-delivery system and design files for the two-photon mosquito holder are available at https://github.com/mcbridelab/Zhao_2020_HumanOdorRepresentation. For the novel analysis pipeline for volumetric antennal-lobe imaging, code is available at https://github.com/rwth-lfb/Zhao_et_al.

---

**Extended Data Fig. 1.**
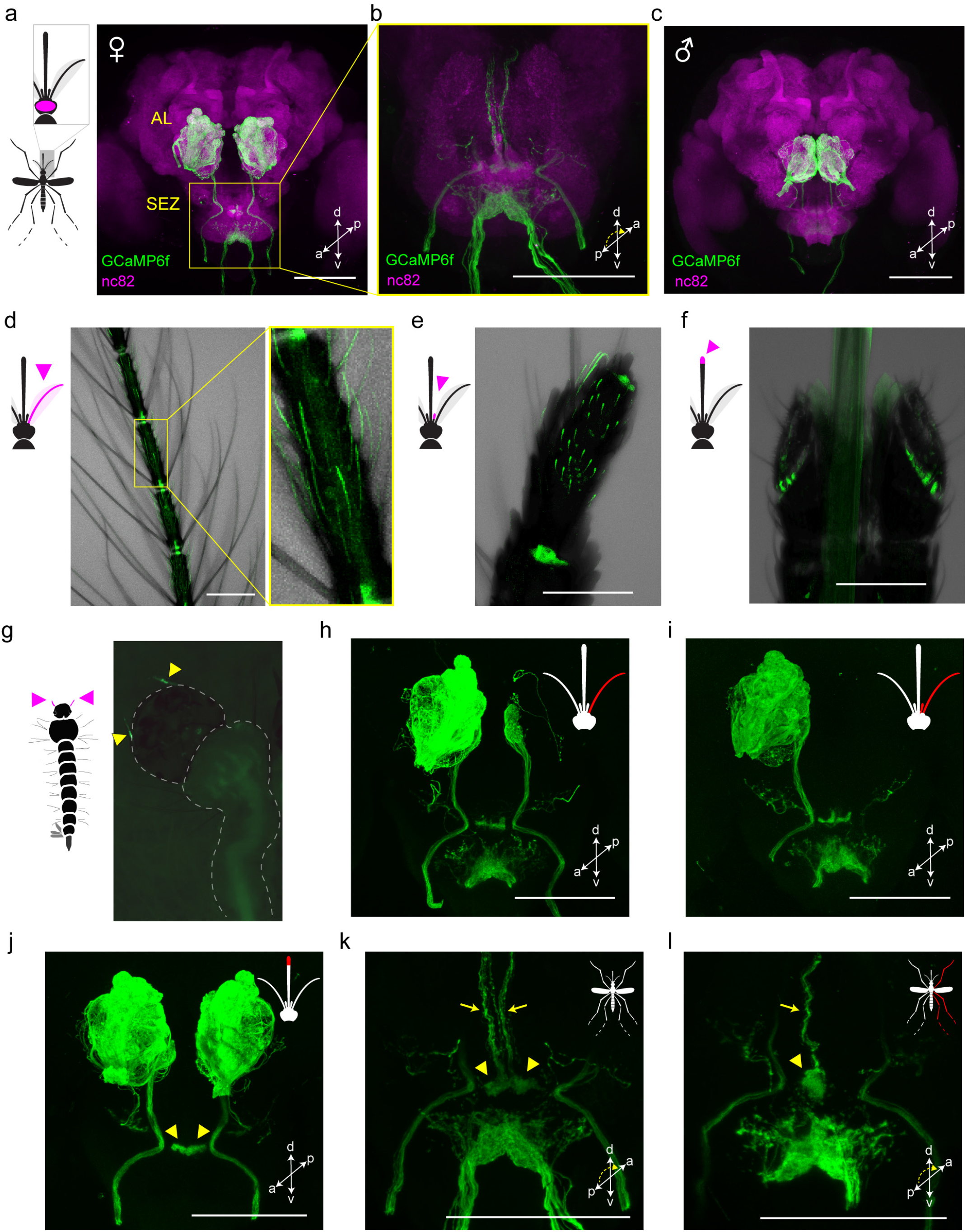
Orco-T2A-QF2-QUAS-GCaMP6f labels chemosensory neurons in peripheral organs that project to the brain. a–c, Antibody staining in female (**a,b**) and male (**c**) brains showing GCaMP6f in sensory neurons that innervate the antennal lobe (AL) and subesophageal zone (SEZ). SEZ in (**b**) is viewed from posterior to better visualize GCaMP6f signal. **d–g**, Intrinsic GCaMP6f fluorescence in sensory neurons of adult female antenna (**d**), maxillary palp (**e**), labella (**f**) and larval antennae (**g**, arrowheads). Transmitted light image overlaid in (**d–f**). **h–l**, Antibody staining in brains of female mosquitoes with severed sensory organs (red in mosquito schematics). Severing right antenna only (**h**), or both right antenna and palp (**i**) led to loss of signal in all ipsilateral glomeruli except two in the posterior-medial region or all ipsilateral glomeruli, respectively. We therefore infer that Orco+ AL glomeruli are innervated by sensory neurons in the ipsilateral antenna (n~32 glomeruli) and palp (n=2 glomeruli). Severing the tip of the proboscis (including the labella) led to loss of signal throughout the ventral SEZ (**j**), consistent with work in *Anopheles gambiae* indicating that Orco+ labellar neurons innervate this region^51^. However, labellum-less animals retained signal in an area of the dorsal SEZ recently termed the subesophageal glomeruli (arrowheads in **j**) (https://www.mosquitobrains.org/). Signal in these glomeruli and corresponding ascending nerves from intact animals (**k**, arrowheads and arrows, respectively) was lost when the ipsilateral legs were severed (**l**), suggesting that *Ae. aegypti* has Orco+ neurons on the legs that project to the SEZ. This is consistent with recent work showing electrophysiological responses to olfactory stimuli on legs^52^. All scale bars 100 μm.

**Extended Data Fig. 2.**
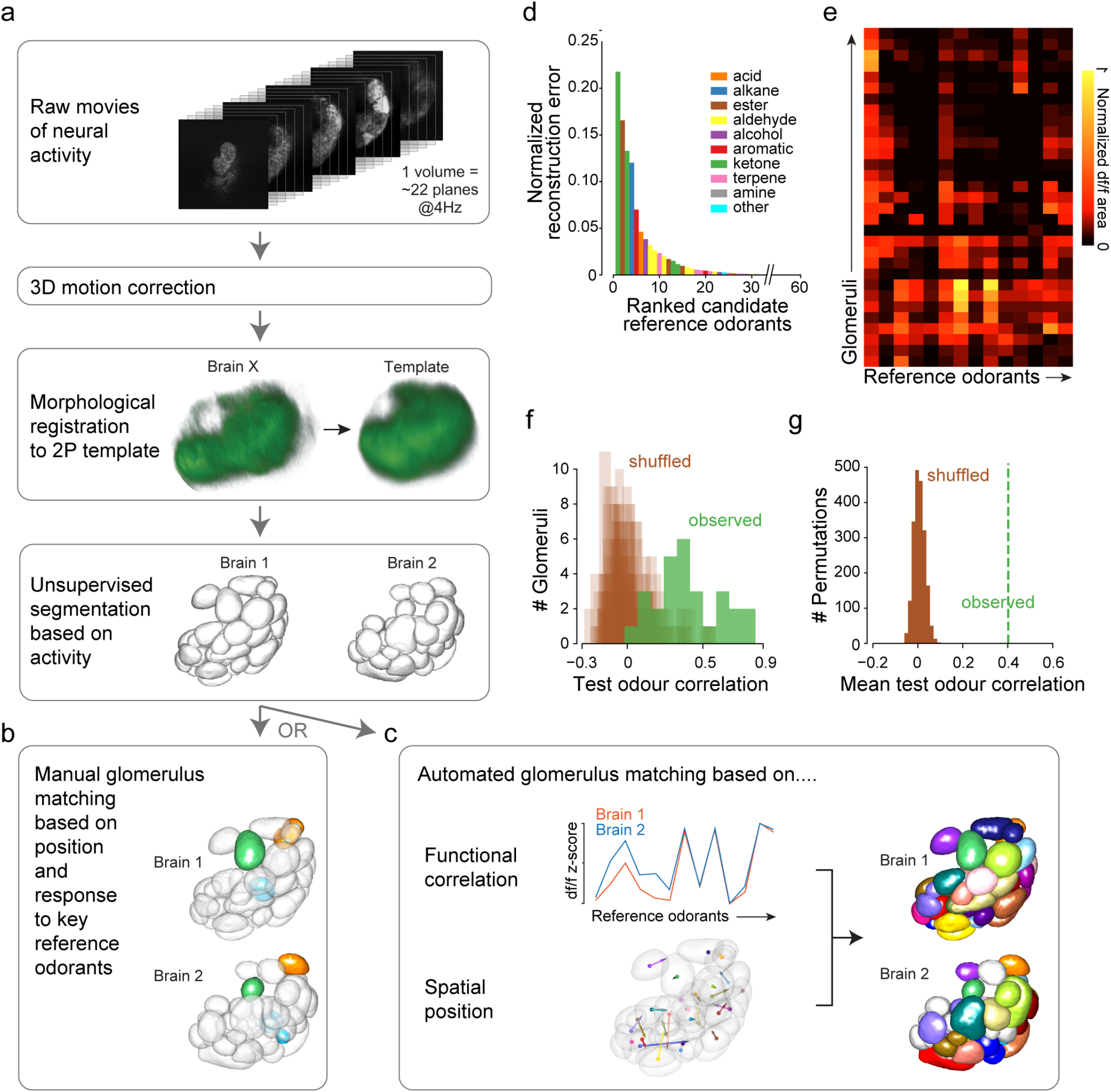
Novel analysis pipeline for volumetric antennal lobe imaging. **a–c**, Pipeline schematic. After registration and unsupervised segmentation of all brains in a given data set (**a**), one brain was chosen as the reference and glomeruli from other brains were matched to those in the reference either manually (**b**) or using an automated pipeline (**c**). Colours in (**b,c**) show matched glomeruli (unmatched in white). Manual matching was performed for analyses focused on only U (cyan), H (green), and A (orange) glomeruli (Fig. 3, 5). Automated matching, which provides a more global picture of activity but is less reliable for the three focal glomeruli, was used for the supplemental analysis in Extended Data Fig. 5b–e. In both cases, matching was based on spatial position and response to 14 reference odorants. **d**, Reference odorants were chosen from among 60 candidates based on their ability to account for a large part of the observed signal variance/neural activity (see Methods). The 10 top-ranked odorants belonged to 8 different chemical classes. **e**, Response of glomeruli from one mosquito to the final 14 reference odorants. **f**, Evaluation of automated glomerulus matching. Glomeruli from 6 brains were matched as in (**c**). We then asked whether the matched glomeruli showed correlated responses to a new set of 13 test odorants. The plot shows the observed distribution of correlation coefficients across n=28 sets of matched glomeruli (green) and 20 shuffled distributions where matches were reassigned at random (brown). Low correlations may be caused by mismatches or general lack of response by a given set of matched glomeruli to the test odorants. **g**, Same as (**f**) except showing the mean of the observed distribution (green line) and the distribution of means from 2000 shuffled datasets.

**Extended Data Fig. 3.**
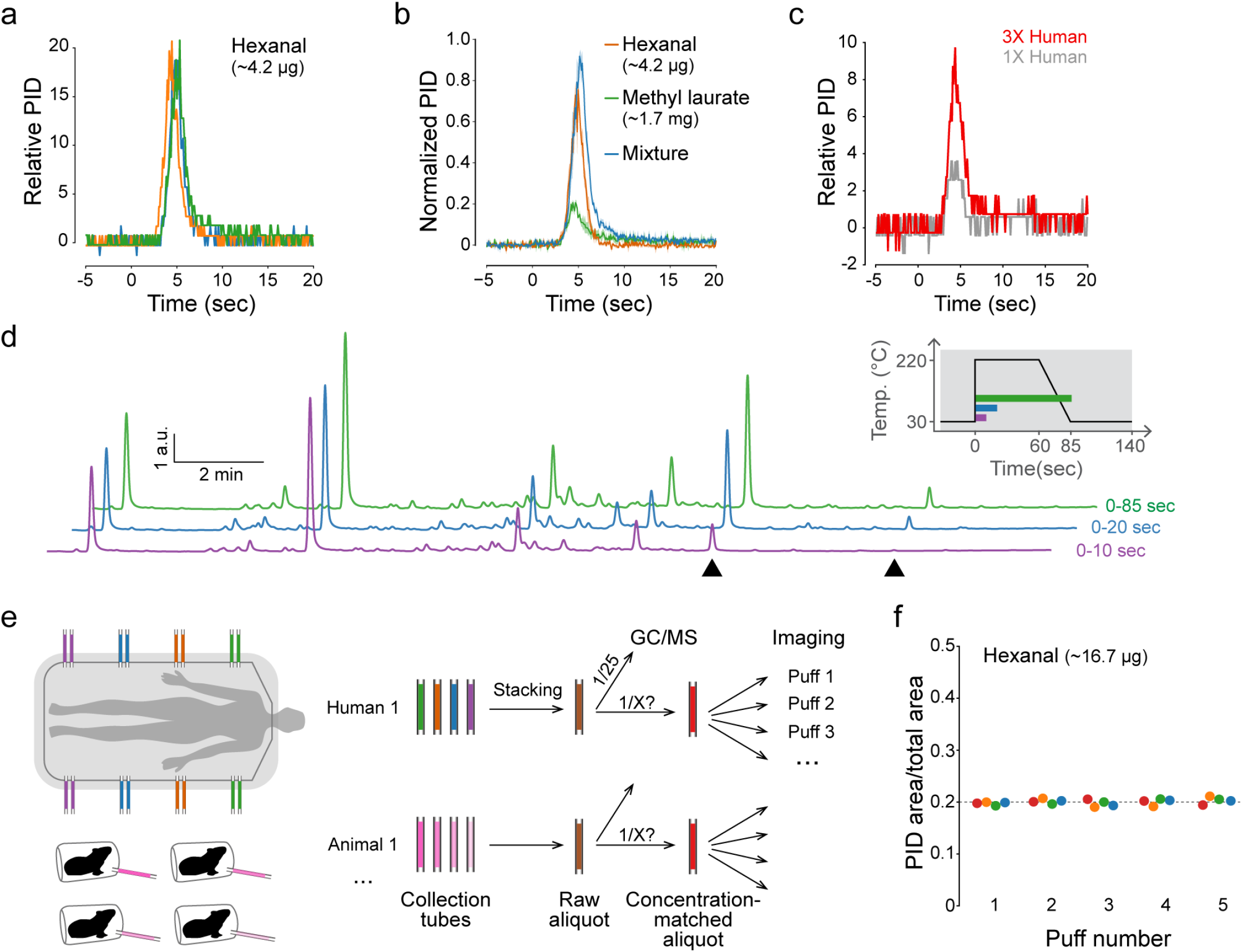
Further characterization of the thermal-desorption odour-delivery system. **a**, Puff shape for hexanal, measured with a photoionization detector (PID) at the location of mosquito antennae in our set-up (n=3 puffs). Time=0 indicates the onset of focusing-trap desorption. It takes ~3 sec for the desorbed odour to reach the mosquito. **b**, Puff shape for hexanal (orange), methyl laurate (green), and their mixture (blue), showing that the temporal dynamics of odour release are similar for odorants with markedly different volatility (n=3 puffs each). **c**. Puff shape of human odour (two doses of the same sample) delivered via thermal desorption and detected using a PID. **d**, GC-MS traces showing the composition of replicate puffs of sheep odour, collected for a period of 10, 20, or 85 seconds following the onset of trap desorption. Inset shows the time over which each puff was collected with respect to focusing-trap temperature. Almost all components were released within the first 10 sec (purple), but arrowheads mark two compounds that took longer to fully desorb. **e**, Schematic of process for pooling (‘stacking’) odour samples and matching their concentrations before use in imaging. We stacked multiple collection tubes from the same individual human subject or multiple individuals of the same animal species to generate a single raw aliquot (brown). We then quantified 1/25th of each raw aliquot via GC-MS in order to make concentration-matched aliquots (red) with the same total odour content (Fig. 2g). **f**, Concentration of five replicate puffs of hexanal delivered from each of four sample tubes (different colours) demonstrating repeatability of the delivered stimulus.

**Extended Data Fig. 4.**
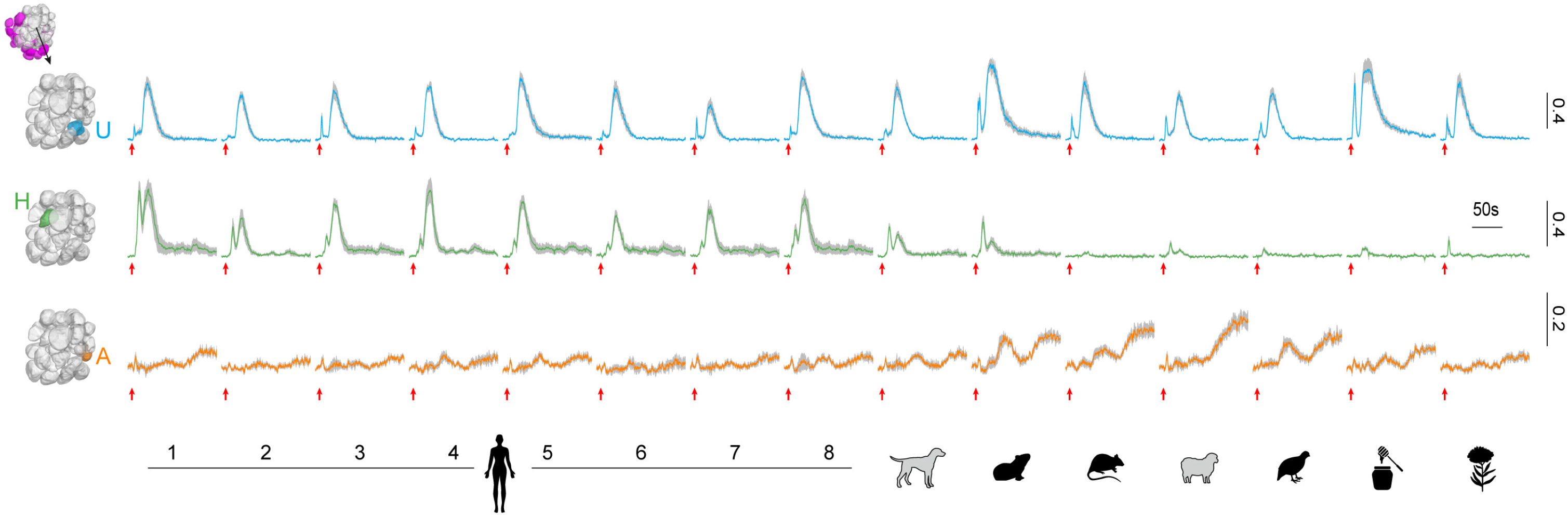
Response of three target glomeruli to diverse odour extracts. Coloured lines and grey shading show mean ± SEM response to 1X concentrations of the given stimuli (n=5 mosquitoes). Red arrows under each activity trace mark the onset of heating. The double peaks observed in some traces may reflect technical aspects of the delivery system (*e.g*. incomplete synchronization of odorant release, Extended Data Fig. 3d). Since synergistic interaction at the OSN level is rare in other insects^17^, we do not expect the asynchrony to greatly impact our findings. We observed neural activity that lasts tens of seconds longer than the stimulus, as well as markedly tonic responses in glomerulus A, for both blends delivered via thermal desorption (this figure) and single odorants or synthetic blends delivered in a more traditional way (Extended Data Fig. 8). Scale bars at right indicate normalized df/f.

**Extended Data Fig. 5.**
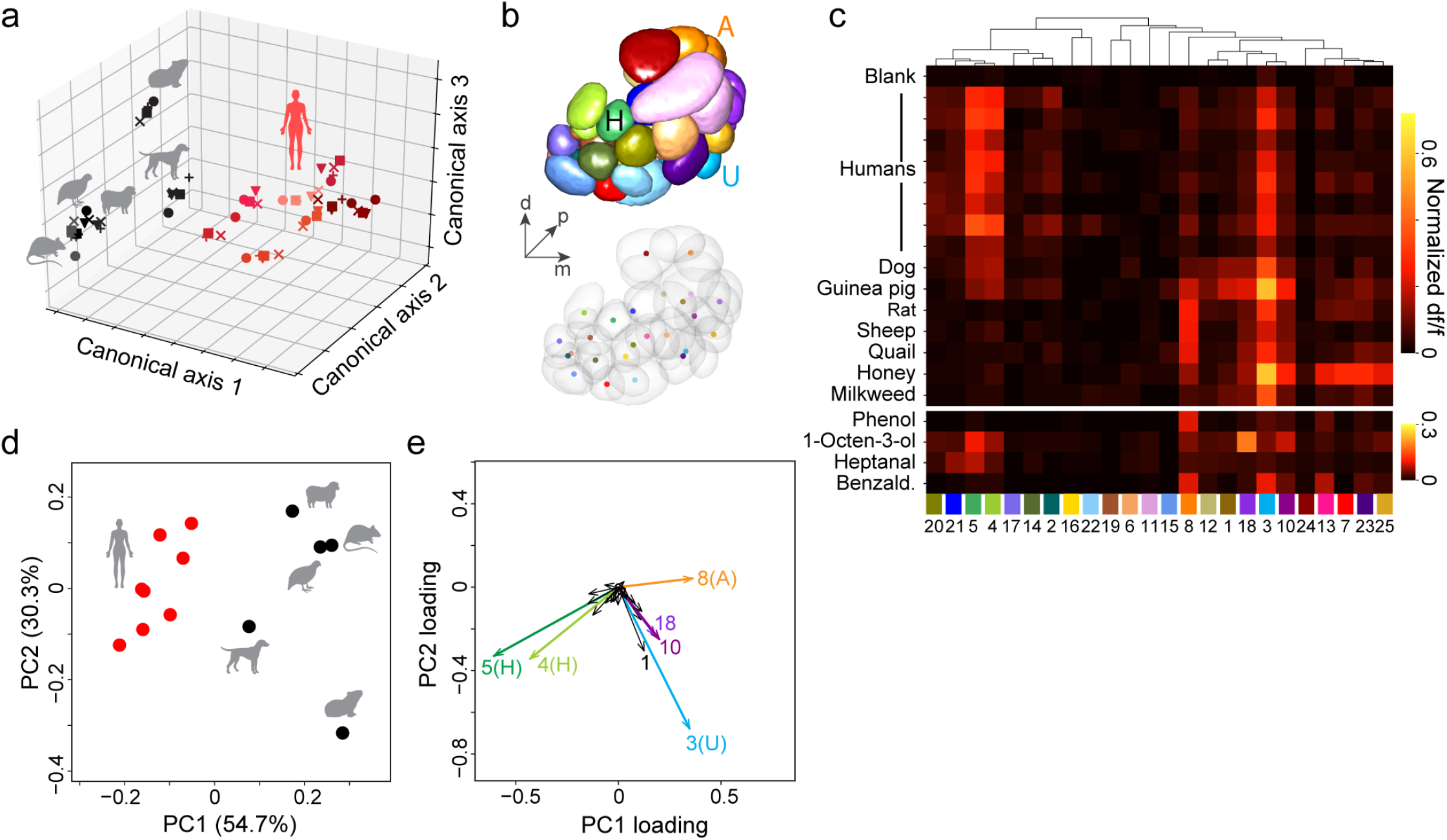
Automated analysis of response to human and animal odours is consistent with targeted analysis of U, H, and A glomeruli. Panels show reanalyses of data presented in Fig. 3h–k. **a**, Human and animal odours were cleanly separated along the first three axes of an across-matrix PCA of unmatched signal clusters from all mosquitoes (see Methods). Symbols denote individual mosquitoes (n=5); shades of red denote odour from different human subjects (n=8). **b–e**, Human and animal odours were also cleanly separated in an analysis of signal clusters matched by the automated algorithm (Extended Data Fig. 2c–g, see Methods). Panel (**b**) shows signal clusters from the segmented antennal lobe of the reference mosquito. Panel (**c**) shows the mean normalized response to odour extracts (top) and select reference odorants (bottom) for those signal clusters that could be matched in the brains of at least 2 of 4 additional mosquito replicates (n=3–5 mosquitoes total per cluster). Panels (**d,e**) show a principal components analysis of mean responses from (**c**). Note that unsupervised segmentation sometimes splits a single glomerulus into two or more signal clusters. To partially address this issue, we merged adjacent clusters with reference-odorant-response correlations above 0.7 prior to matching (see Methods). ‘Merged’ clusters from the reference mosquito were assigned the same colour in (**b**) *(e.g*. the two dark-orange A clusters). The adjacent green and lime-green clusters in (**b**) did not quite meet the threshold for merging, but nevertheless show correlated responses (**c,e**) and likely both correspond to the H glomerulus. Responses to human and animal odours were cleanly separated (**d**) based primarily on activity in four major signal clusters (**e**) corresponding to H (#4 and 5), A (#8), and U (#3) glomeruli. A few other clusters also had substantial loadings, including the purplish clusters #10 and 18, which are also spatially adjacent and likely correspond to the same glomerulus. This glomerulus was obscured in the manual analysis because it is just posterior to U and has partially correlated, more or less universal responses to vertebrate odours. However, unlike U, it is strongly activated by the reference odorant 1-octen-3-ol (**c**), a known ligand of palp neurons that project to this posterior-medial region of the antennal lobe.

**Extended Data Fig. 6.**
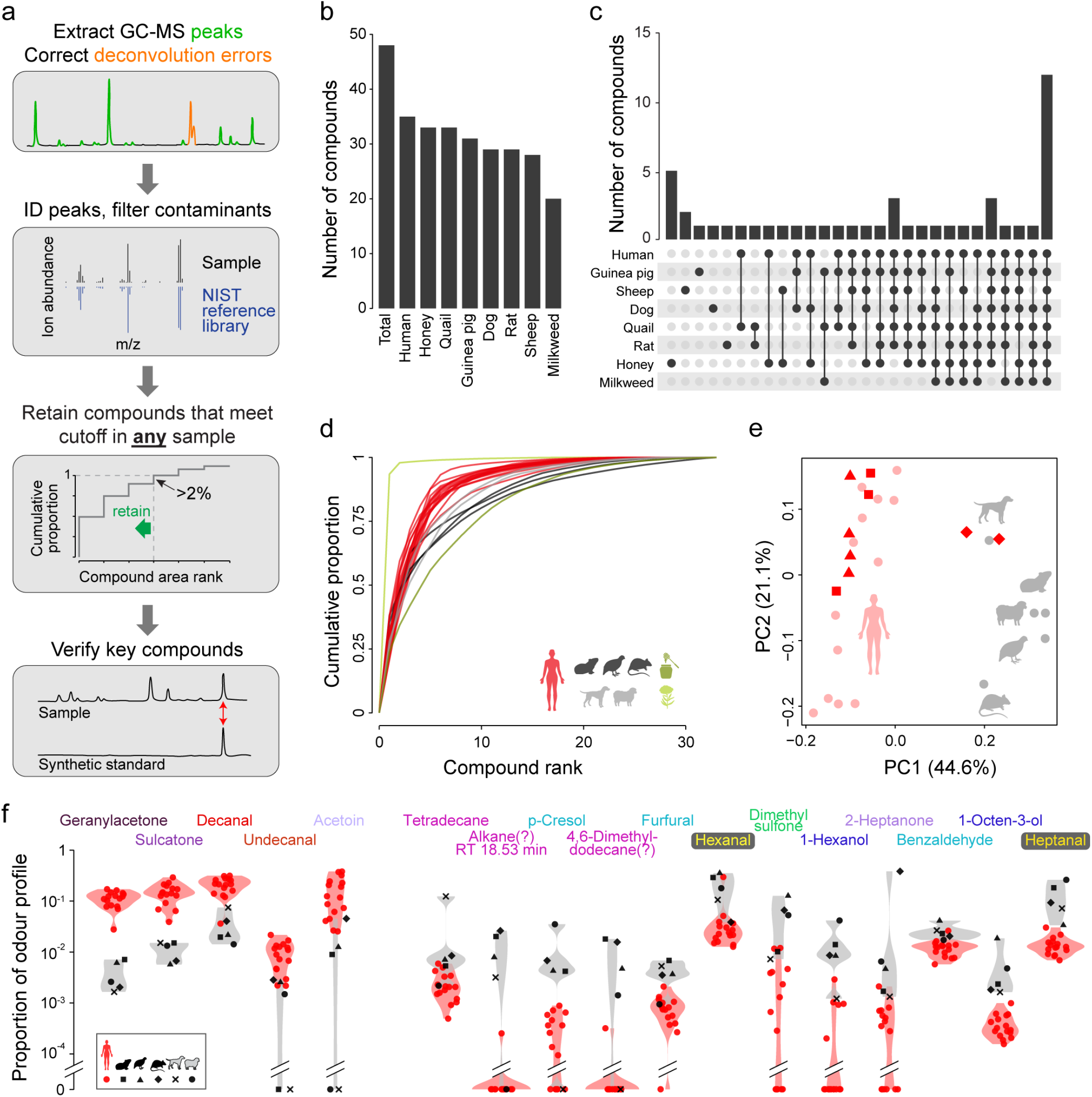
Quantitative analysis of human odour and animal odours. **a,** Analysis pipeline for GC-MS data. **b**, Number of compounds found in each species/stimulus. **c**, Number of compounds found in the given combination of species/stimuli. **d**, Cumulative distribution of odorants in each odour profile. **e**, Unscaled principal components analysis of human and animal odour profiles similar to Fig. 4b but including 2–4 replicate odour extractions for three of the human subjects (dark red, different symbols denote different subjects). **f**, Violin plots showing on a log scale the relative abundance of odorants that passed the significance threshold in Fig. 4e.

**Extended Data Fig. 7.**
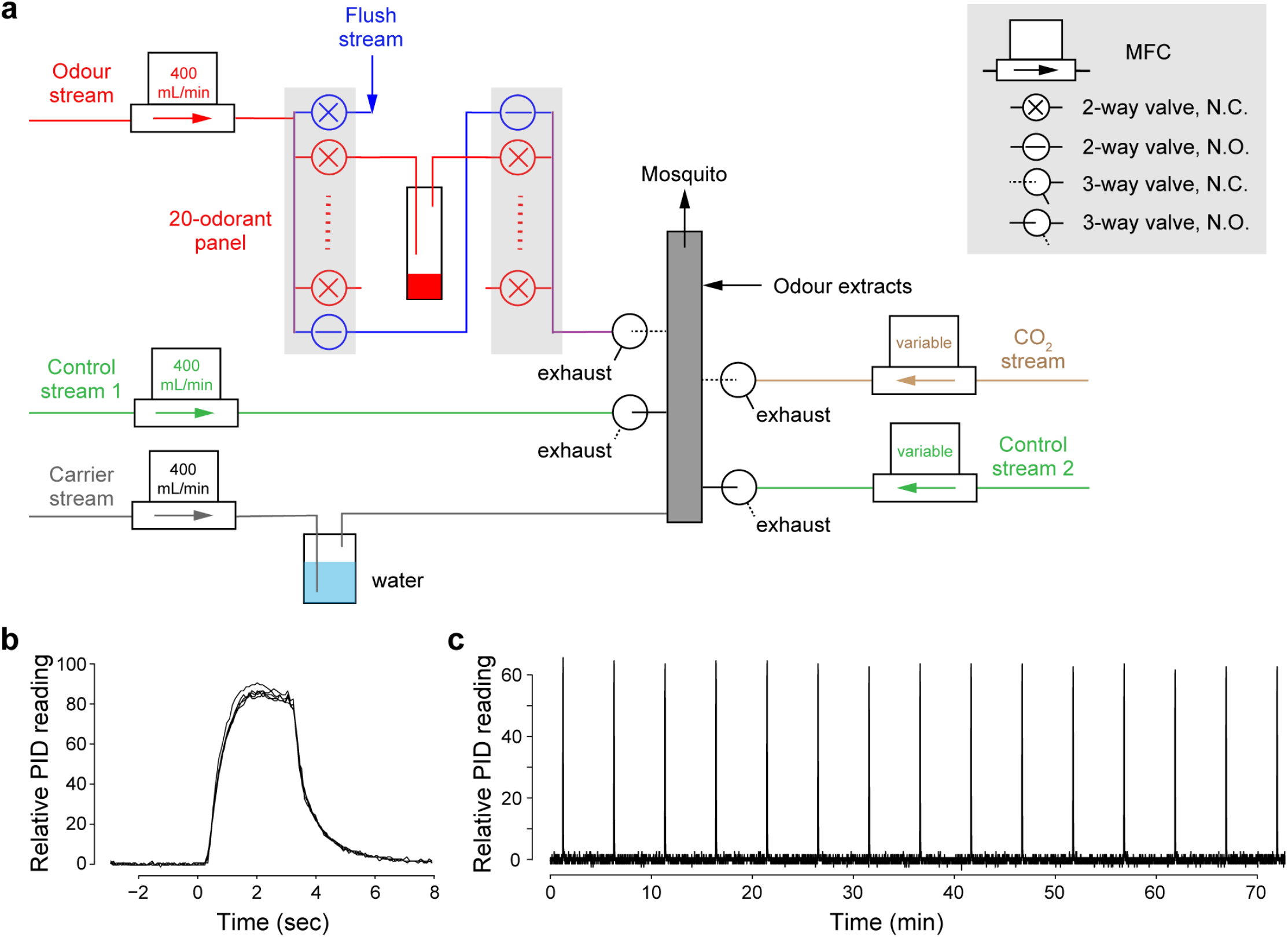
Design and characterization of the single-odorant delivery system. **a,** Design schematic. Filtered air is split into 5 streams, each regulated by a mass flow controller (MFC). The humidified carrier stream (grey) flows continuously through the mixing manifold (grey box) to the mosquito. Normally, the two control streams (green) are also flowing through 3-way valves to the manifold. Synthetic odorants and CO_2_ are delivered through the odour stream (red) and CO_2_ stream (brown), respectively. The odour stream has 20 odour channels (red) plus a bypass (blue). To puff the odorant in a given vial, the bypass closes, 2-way valves flanking the odour vial open, and the headspace of the odour vial is carried by the odour stream to a 3-way valve that diverts the stream from exhaust to the mixing manifold with a delay. Meanwhile, control stream 1 is diverted to exhaust to maintain a constant flow rate. When delivering CO_2_, the CO_2_ stream (fed by a carbogen tank) is similarly diverted to the mixing manifold and offset by control stream 2. The high-flow flush (blue, 2000 mL/min) opens between odour puffs to remove residual odorant from the system. Output of the thermal-desorption system used to deliver complex odours also joins the final mixing manifold. MFC, mass-flow controller; N.C., normally closed; N.O., normally open. See Methods for more detail. **b**, Shape of odour puffs delivered by the system, featuring fast rise/decay and stable peak height. Five replicate 3-sec puffs of 2-heptanone (10^-2^ in paraffin oil) were aligned to the command onset (time=0). **c**, Long-term stability of odour puffs delivered by the system. A 3-sec puff of 2-heptanone (10^-2^) was delivered every 5 min for 75 min.

**Extended Data Fig. 8.**
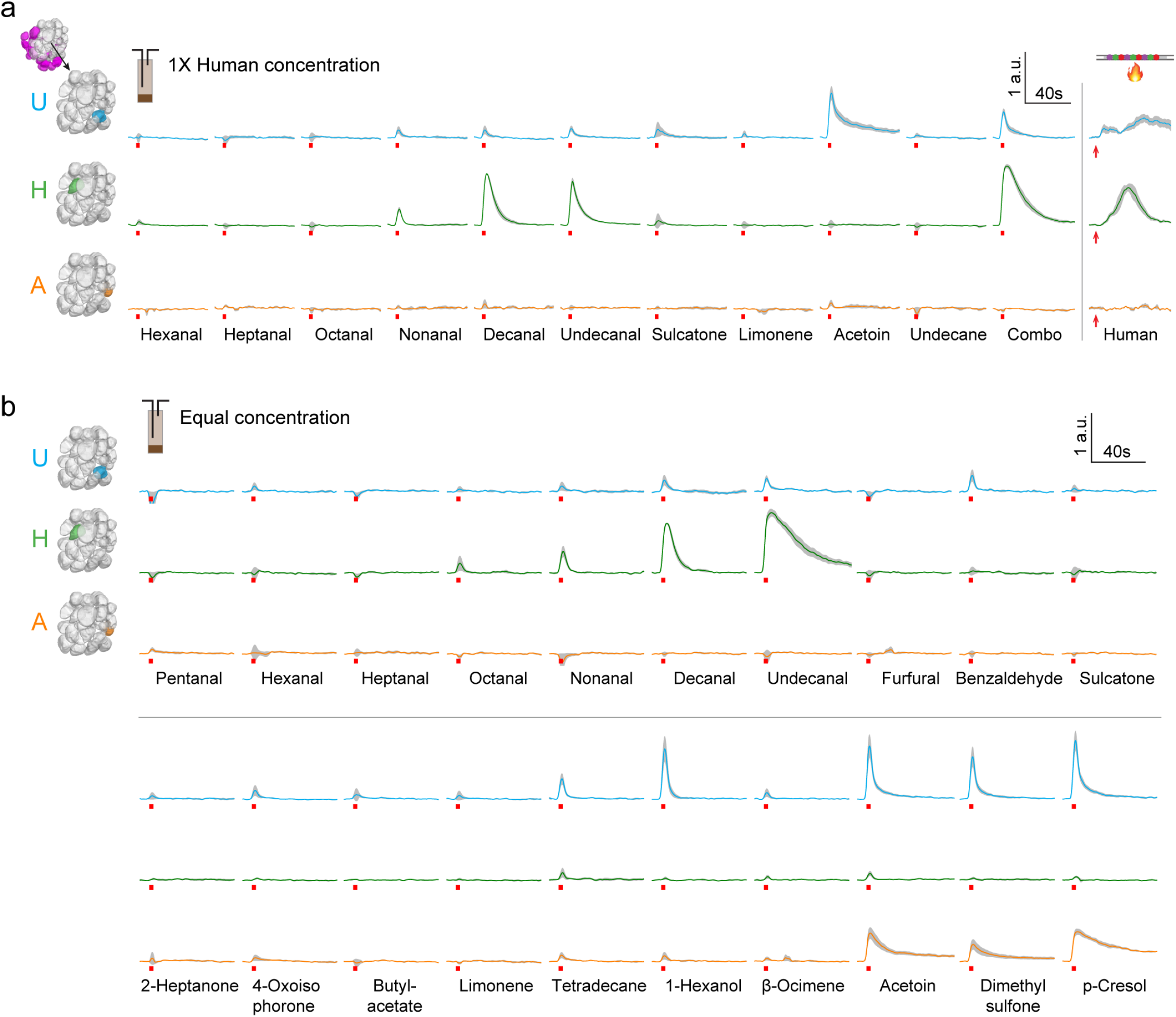
Response of three target glomeruli to single odorants. **a,** Response to major components of human odour delivered at their respective concentrations in a 1X human sample. Combo is a mix of all the individual components except acetoin. Response to a thermally desorbed 1X human-odour puff is shown at far right. **b**, Response to individual odorants delivered at equal vapour-phase concentration. Exceptions were dimethyl sulfone and p-cresol, which were delivered at approximately half the target concentration due to their low volatility and solubility in the chosen solvent (Fig. 5f). In both (**a**) and (**b**), coloured lines and grey shading indicate mean ± SEM for n=4–5 mosquitoes. Red bars mark the timing of the 3-sec puff for each synthetic odorant or combo. Red arrow for human extract in (**a**) marks the onset of focusing-trap desorption. Note that glomerular responses to single components are often prolonged, lasting well beyond the 3 sec stimulus. This is consistent with recent single-sensillum recordings that found a prolonged response by olfactory sensory neurons to certain odorants, including aldehydes^82^. Arbitrary units (a.u.) are df/f normalized within mosquitoes to the response of glomerulus H to decanal delivered at 1X human concentration (**a**) or ‘equal’ concentration (**b**).

